# Synergistic regulation of bZIP53 and dimerizing partners results in abnormal seed phenotype in Arabidopsis: Use of a designed dominant negative protein A-ZIP53

**DOI:** 10.1101/2021.07.20.453159

**Authors:** Prateek Jain, Vikas Rishi

**Affiliations:** National Agri-Food Biotechnology Institute, Knowledge City, Sector 81, Mohali, Punjab 140306, India

## Abstract

In Arabidopsis, basic leucine zipper (bZIP) family of transcription factors (TFs) are key proteins to regulate the expression of seed maturation (MAT) genes. bZIPs are functionally redundant and their DNA-binding activity is dependent on dimerization partner. The intervention of loss of function mutation is inadequate to understand and regulate the redundant behavior of TFs and one such example is bZIP53, which is known as a key regulator of seed maturation phenomena. Here, to examine the consequences of hindering the function of bZIP53 and its known and unknown heterodimerizing partners, a transgenic *Arabidopsis* constitutively expressing a novel dominant negative (DN) protein A-ZIP53 was raised. Transgenic plants demonstrated a delayed growth and retarded seed phenotype. The *in vivo* inhibition of DNA binding of bZIP53, bZIP10, and bZIP25 to the G-box demonstrated the efficacy of A-ZIP53 protein. In first generation, majority of plants failed to survive beyond four weeks suggesting a pleiotropic nature of bZIP53. Plants expressing *A-ZIP53* have small flower, shorter siliques, and small-seeded phenotype. RNA seq analysis of the transgenic lines revealed the reduced expression of target genes of bZIP53 and its heterodimerizing partners. Furthermore, immunoprecipitation followed by mass spectrometry (IP-MS/MS) of transgenic plants helped to identify the additional heterodimerizing partners of the A-ZIP53. The interactions were subsequently confirmed with the transient transfection experiments. Unlike other gene knock out technologies, DN protein can inhibit the function of members of the same group of bZIP TFs.

## Introduction

Gene expression is controlled by dynamic interaction between various biological components including hormones, small RNAs, transcription factor (TFs), epigenetic modification etc. These interactions are spatial or temporal in which DNA binding TFs are master regulators. TFs regulate the function of a target gene in an additive fashion or cooperative manner by the homo- or heterotypic interactions. In *Arabidopsis*, these dimeric interactions regulate several process including seed development and maturation, which is well studied and explored (Alonso et al. 2009; Santos-Mendoza et al. 2008).

Seed maturation is a fine-tuned process, which occurs only in angiosperm. This process collectively occurs in different part of seed, which ultimately contributes to seed quality (Vicente-Carbajosa and Carbonero 2004; Kumar et al. 2018). It is divided into different phases in which early and mid-phase are dominated by action of ABA and in late phase ABA level decreases followed by the synthesis of Late embryogenesis abundant (LEA) protein (Tunnacliffe and Wise 2007). Seed maturation is a complex process and the intervention of high throughput genome and transcriptome technologies have helped to identify target genes (Sreenivasulu and Wobus 2013). Maturation phase is controlled singly or in combination via seed storage protein genes (SSP) such as, *ABI3*, *FUS3*, and *LEC1* (Braybrook et al. 2006; Jakoby et al. 2002; Parcy et al. 1994; Santos-Mendoza et al. 2008). Mutation in these genes result in severely affected seed phenotype with reduced content of seed storage proteins (To et al. 2006). Besides defects in seed development, mutants display other pleiotropic effects like chlorophyll accumulation in dry seeds, desiccation intolerance, defected cotyledon identity, and others (Bensmihen et al. 2005; Boyes et al. 2001; Keith et al. 1994; Kroj et al. 2003; Meinke et al. 1998; Meinke et al. 1994; Parcy et al. 1994; Stone et al. 2001). Although, previous studies have shown the involvement of these genes in seed development, but the mechanism regarding their interaction and regulation is not well defined (To et al. 2006). The expression of Maturation associated (MAT) genes are under the tight control of several cis regulatory element including ACGT elements, RY (CATGCA), AACA, and CTTT motifs (Vicente-Carbajosa and Carbonero 2004), which are the binding site for various transcription factors including B3, MYB, and basic leucine zipper (bZIP) (Santos-Mendoza et al. 2008).

bZIPs are the eukaryotic class of TFs that bind to DNA in a sequence specific manner. It is a bipartite structure in which N-terminal act as a DNA binding domain while dimerization is done by C-terminal. These TFs bind to their cognate site as a dimer. These dimeric interactions are dynamic and responsible for the homo- or heterotypic interactions (Acharya et al. 2006; Amoutzias et al. 2008; Landschulz et al. 1988). Structurally, bZIP dimerization depends on amino acid present at the ‘a’, ‘d’, ‘e’, and ‘g’ positions in heptad, which defines the affinity and specificity of the dimeric interactions (Deppmann et al. 2006; Vinson et al. 1993). These dimeric interactions are responsible for redundant behavior of bZIP TFs (Alonso et al. 2009; Dietrich et al. 2011).

In *Arabidopsis thaliana*, sequence similarity have been used to predict the number and possible dimerization partner of bZIPs (Deppmann et al. 2006; Jakoby et al. 2002). They can form a network, which regulate the myriad of biological processes. As reported earlier, bZIP of Group C and S (Jakoby et al. 2002) can form C/S1 network and participate in seed maturation (Alonso et al. 2009), energy homeostasis (Baena-González et al. 2007), amino acid metabolism (Dietrich et al. 2011; Weltmeier et al. 2006; Weltmeier et al. 2009), and salt stress (Hartmann et al. 2015). Several bZIPs like ABI–5, bZIP67 (Belmonte et al. 2013), bZIP15, and bZIP72 (Le et al. 2010) are known to be involved in seed maturation and among these, bZIP53 has been reported as a key regulator of maturation associated genes (MAT) (Alonso et al. 2009). bZIP53 with its heterodimerizing partners bZIP10 or bZIP25 is involved in the regulation of MAT genes expression (Alonso et al. 2009). On the other hand, bZIP53|bZIP1 heterodimer has been reported in salt stress (Hartmann et al. 2015). In general, the dimerizing partner selection is responsible for the pleiotropic and redundant behavior of target bZIP TFs.

Recently, we have reported a novel dominant negative (DN) protein approach to regulate the dimeric interaction of target bZIP TFs involved in seed maturation (Jain et al. 2018; Jain et al. 2017). The efficacy of novel protein A-ZIP53 and its derivatives against wild-type bZIP53 and its heterodimerizing partners were shown *in vitro* and *in vivo* (Jain et al. 2018; Jain et al. 2017). Here, we have generated transgenic *Arabidopsis* expressing DN protein A-ZIP53 under the constitutive promoter and examined the effects on the growth and development. Transgenic lines have a differential growth pattern and abnormal seed phenotype. The effect of A-ZIP53 on the seed maturation was confirmed with the down regulation of seed-specific genes. RNA-seq analaysis helped us to understand the effect of A-ZIP53 expression on the overall expression profile in plants. Furthermore, the immunoprecipitation followed by mass spectrometry (IP-MS) was used to identify the additional heterodimerizing bZIP partners of A-ZIP53. The transient transfection assay confirmed the heterodimeric interaction between the A-ZIP53 and new immunoprecipitated bZIPs. Therefore, we propose the efficacy of the novel designed DN protein *A-ZIP53* against target bZIPs and dimerizing partners. The DN protein can be used to regulate the redundant behavior of the target protein and to predict the unknown dimerizing partners. Ultimately, it will help to understand and regulate the redundant behavior of target bZIP TF.

## Results

### A-ZIP53 transgene expression causes delayed growth phenotype in *Arabidopsis*

In a previous study, we have biochemically characterized dominant negative (DN) protein A-ZIP53 and its derivatives. The efficacies of designed proteins were demonstrated *in vitro* and *ex vivo* (Jain et al. 2018; Jain et al. 2017). A-ZIP53 is designed to specifically target bZIP53 and its known heterodimeric partners and inhibit their DNA binding thus down-regulating associated genes. Earlier in gel shift experiments and transient transfection studies, we have shown the ability of A-ZIP53 in inhibiting DNA binding activities of bZIP53 and its known dimerizing partners i.e., bZIP10 and bZIP25 (Alonso et al. 2009; Hartmann et al. 2015; Jain et al. 2018; Jain et al. 2017). The interaction between bZIP53 and other bZIPs like bZIP10, bZIP25, bZIP63, and bZIP14 were also confirmed using yeast two-hybrid system and applying transient transfections in *Arabidopsis* protoplast (Ehlert et al. 2006; Weltmeier et al. 2006; Weltmeier et al. 2009), suggesting a pleotropic behavior of bZIP53 with a possible role(s) in different physiological and metabolic pathways (Alonso et al. 2009; Hartmann et al. 2015). To understand the consequences of inhibiting DNA binding activities of bZIP53 and its heterodimerizing partners in seed maturation, A-ZIP53 was expressed under a constitutive promoter (p35S: A-ZIP53) in the wild type *Arabidopsis* (Col-0 background) (Figure 1A).

**Figure1.**
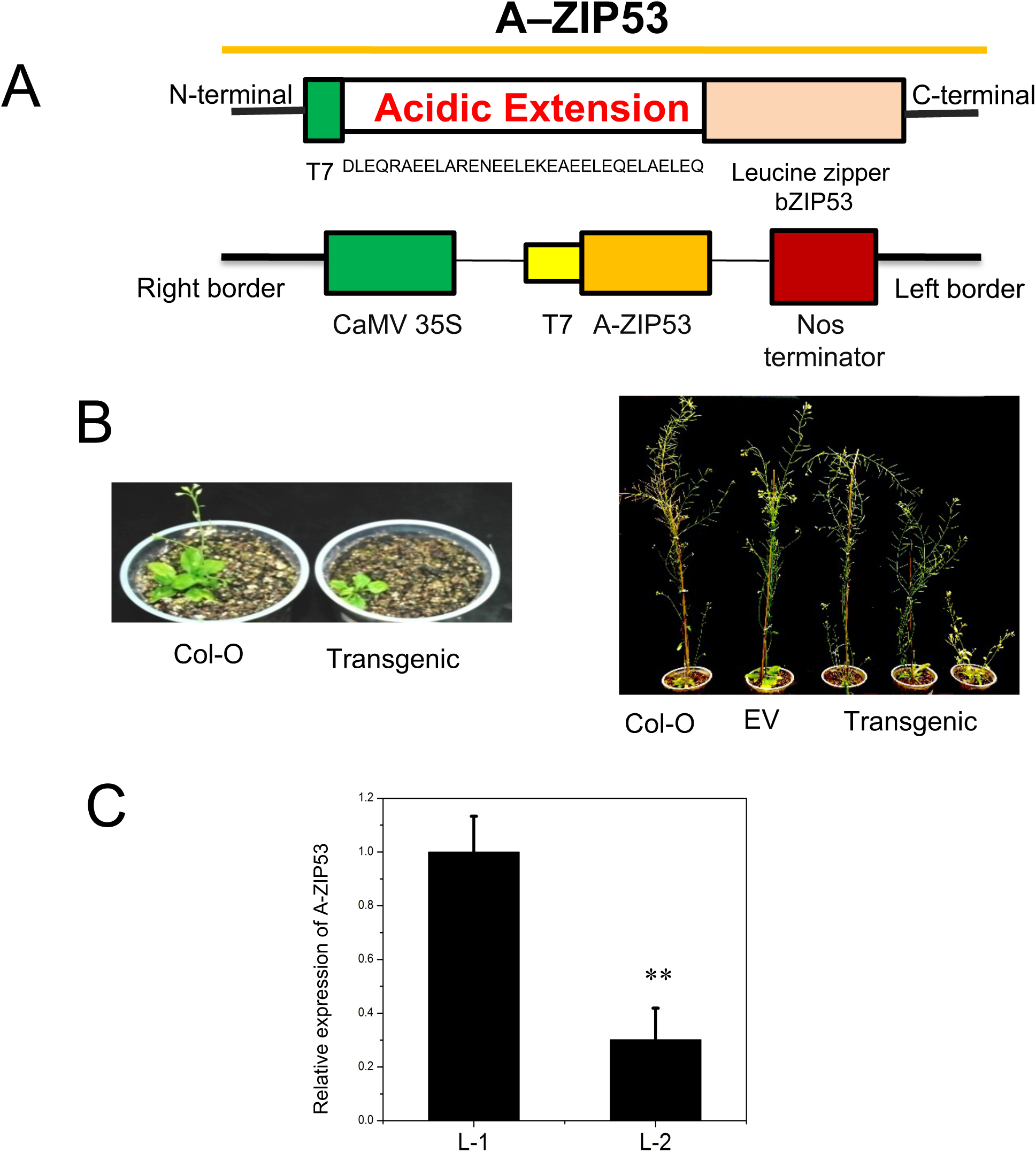
Constitutive expression of A-ZIP53 in wild-type *Arabidopsis thaliana* **(A)** Schematic of Pro35S: A–ZIP53 construct as described in Materials and method. **(B)** Phenotype alternation and abnormal growth pattern of 12 weeks old transgenic *Arabidopsis* of T -1 generation. **(C)** Differential expression of *A-ZIP53* in different lines of transgenic. Error bar represent (±) mean and S.D of three biological replicates. The level of significance was calculated using oneway ANOVA. ** level of significance (P < .01). Error bar indicates mean and S.D. of three biological replicates.

A-ZIP53 expressing transgenic plants were analyzed for growth and other physiological parameters. Most of transgenic plants showed altered phenotypes including retarded growth, dwarfism, and late flowering compared to wild type. A positive correlation was observed between expression levels of A-ZIP53 as confirmed by qRT-PCR and severity of the phenotype (Figure 1B, C).

Because of the apparent differential expression of A-ZIP53 and corresponding phenotype, independent transformants of *Arabidopsis* were identified and carried for the next generation. The chosen lines were probed for A-ZIP53 by Western blots and lines with a high expression were selected (Figure 2A). Unlike wildtype, A-ZIP53 expressing transgenic plants showed delayed growth phenotype (Figure 2 B and 2C). Plants had stunted growth, small sized flower, shorter silique, and small seeds. Number of seed in the individual silique were 8 -10 compared to >40 in the wild type (Figure 2D). These results and earlier studies suggest severe phenotype in A-ZIP53 expressing plants may be due to inhibition of DNA binding activity of bZIP53 and its known and unknown partners (Hartmann et al. 2015; Alonso et al. 2009).

**Figure2.**
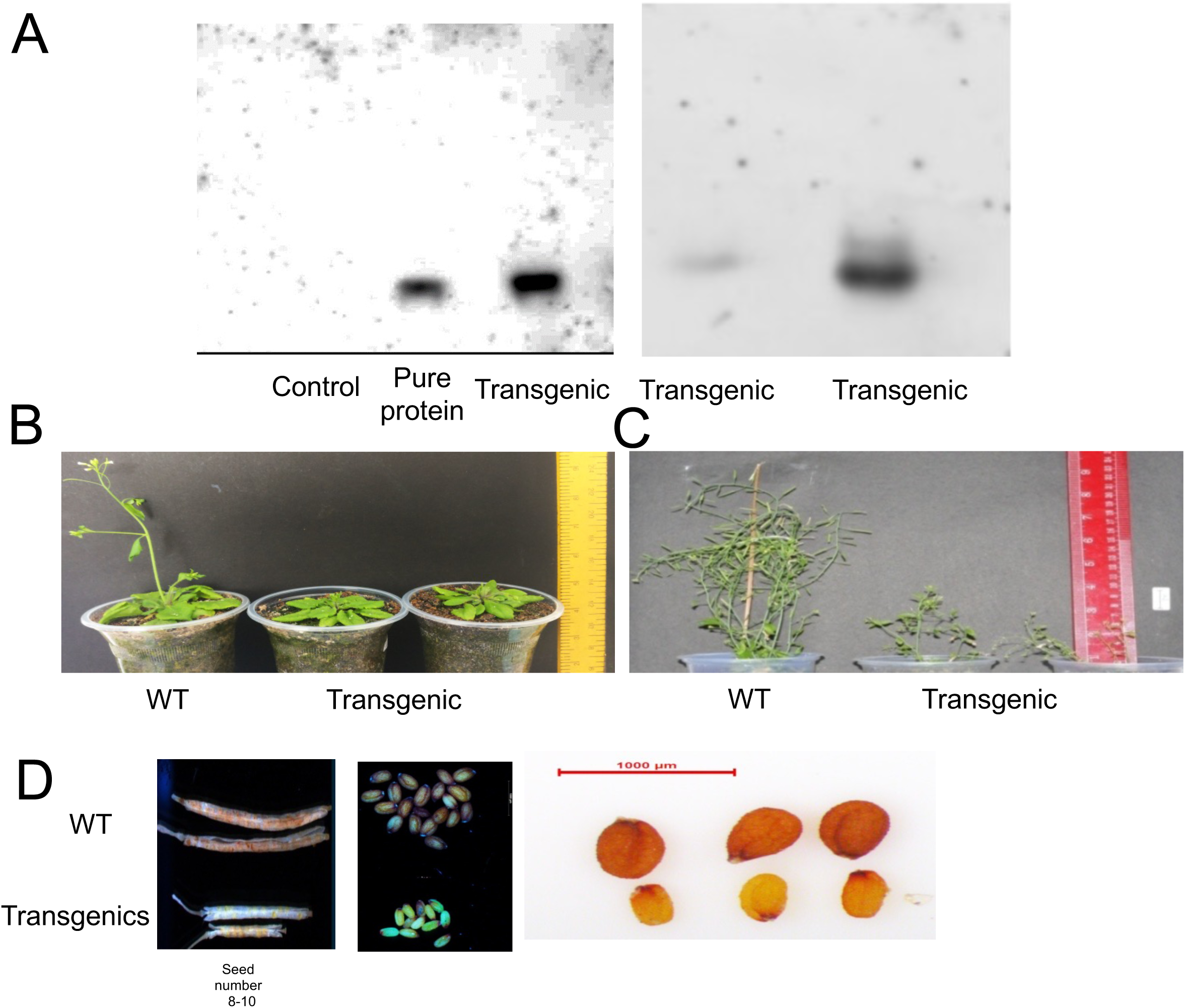
**(A)** Western blot confirms the expression of A-ZIP53 in five days old seedling. **(B)** Phenotypic variation in the growth of four weeks old transgenic compared to wild type. **(C)** Differences in the growth of six weeks old transgenic and wild-type. **(D)** Variation in the silique and seeds of wild type and transgenic.

### A-ZIP53 inhibits the DNA binding of *bZIP53*, *bZIP10*, and *bZIP25*

bZIP53 plays a central role in regulating seed maturation genes and ectopic expression of *bZIP53* resulted in the activation of seed-specific genes (Alonso et al. 2009). bZIP53 binds to DNA as homodimer or as heterodimer with bZIP10, and bZIP25. In earlier we have quantified these interactions using biophysical techniques and gel shift experiments (Jain et al. 2018; Jain et al. 2017). To investigate the ability of A-ZIP53 in inhibiting the functions of target bZIP53 and its interacting partners, transient transfections were performed using protoplast derived from BY-2 tobacco cell line. For transfection assays, DNA construct coding A-ZIP53, bZIP53, bZIP10, and bZIP25 were co-transfected with the reporter plasmid (GUS expression under 2S2 promoter) and control plasmid (35S:NAN). Reduced GUS signals were observed when cells were co-transfected with A-ZIP53 plasmid and showed dose-dependence. Signals were normalized to those of 35S:NAN control plasmid (Figure 3A). Furthermore, GUS signal increased significantly when cells were co-transfected with bZIP53, bZIP10, and bZIP25 suggesting heterotypic interaction between bzip53, bzip10, and bzip25. Interestingly, when cells were co-transfected with A-ZIP53 under above mentioned conditions, GUS signals decreased significantly strongly suggesting that A-ZIP53 can inhibit the activities of all three bZIP TFs (Figure 3B). These results confirmed the *in vivo* efficacy of the A-ZIP53 against target bZIP TFs.

**Figure 3.**
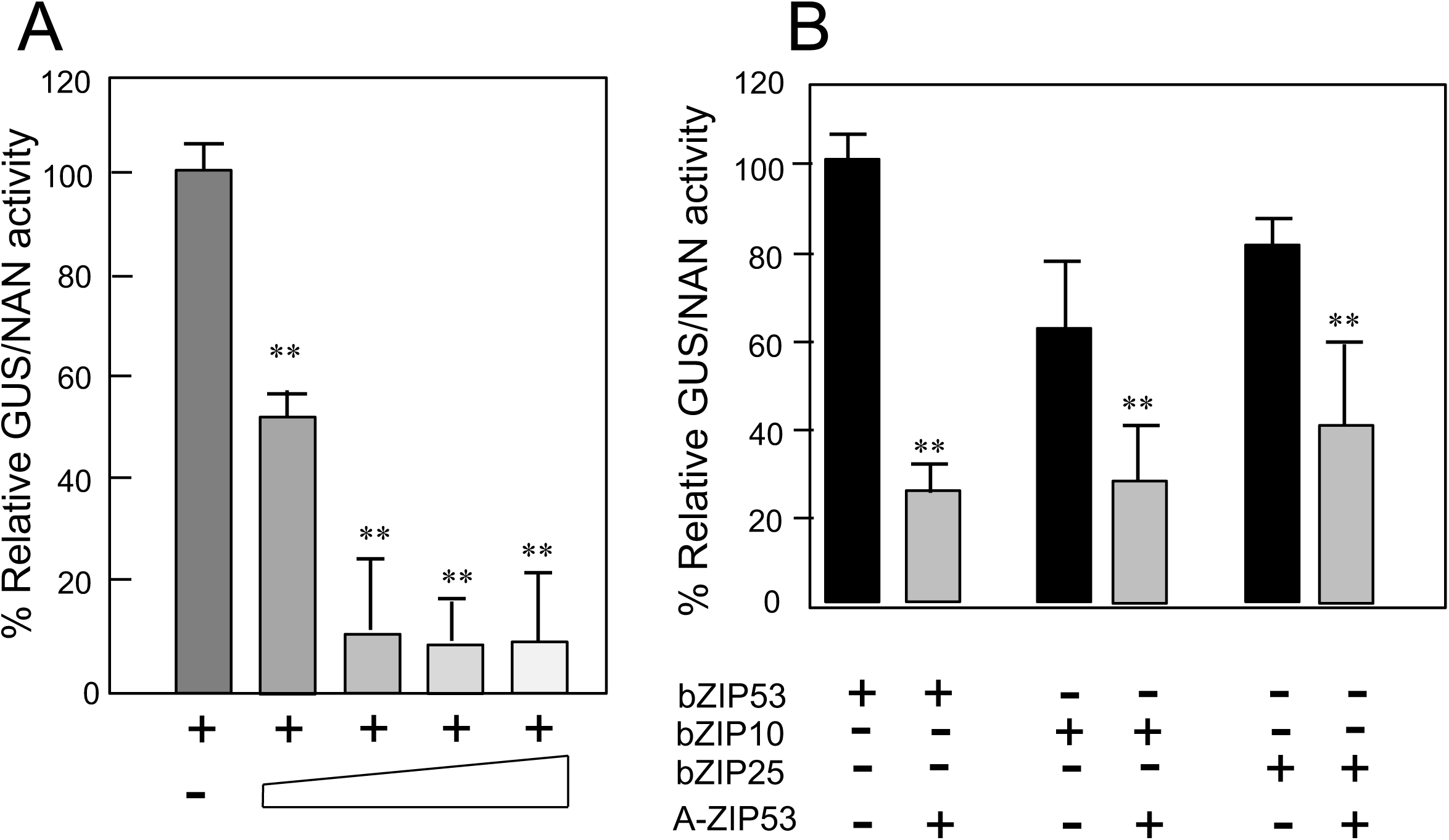
*A-ZIP53* hinders the DNA binding activity of target bZIPs in the transient transfection assay using *Arabidopsis* protoplast. **(A) Dose dependent Assay:** 9 µg of bZIP53 and the indicated increasing molar eq of A-ZIP53 plasmids were co transformed into the protoplast. The y axis defined as relative GUS/NAN activity. **(B)** BY-2 cell line protoplasts were transformed with 9 µg of bZIPs (53, 10, and 25) with 2 molar eq of A-ZIP53. 35S:NAN used as an internal control plasmid (1 µg). The Y axis represented as relative GUS/NAN activity (Alonso et al., 2009). Error bar represent (±) mean and S.D of three replicates from three independent transfection. The level of significance was calculated using oneway ANOVA. *** level of significance (P < .01). Error bar indicates mean and S.D. of three biological replicates.

### *in-silico* promoter analysis of target genes

Protein binding microarray (Weirauch et al. 2014), DAP-seq (O’Malley et al. 2016), Bimolecular Fluorescent Complementation (BiFC) (Llorca et al. 2015), ChIP, and bZIP over expression data used to predict target genes of bZIP TFs potentially participate in seed development and maturation (Weltmeier et al. 2009; Alonso et al. 2009; Weltmeier et al. 2006). Seven genes comprising cruciferin (*CRU*), asparagine synthase1 (*ASN1*), cruciferina (*CRA*), hydroxysteroid dehydrogenase 1(*HSD1*), seed storage albumin (*2S2*), Proline dehydrogenase (*ProDH*), and late embryogenesis accumulating 76 (*LEA76*), (Supplementary table S1) (Alonso et al. 2009)shortlisted for promoter analysis to mark the presence of G-box (<u>ACGTG</u>) (Table I), a potential known binding site for bZIP10 and bZIP25(Lara et al. 2003). **Interestingly,** promoter of target genes also possesses the DNA binding sites for other bZIP TF including bZIP39 (<u>ACGTG</u>) that suggests a possible interaction and functional synergism between two bZIPs.

**Table I.** **–** Putative heterodimeric target bZIPs were revealed using IP-MS against the T7 antibody in the protein soup of A-ZIP53 expressing immature silique and seeds and proteins separated on 15 % SDS PAGE.

|  | Transgenic |  | A-ZIP53 protein incubated with whole protein extract from wild-type Arabidopsis |
| --- | --- | --- | --- |
|  | Immature siliques and seeds | Leaves |  |
| Immuno-precipitated bZIPs | 29, 14, 33, 17, 33, 69, 10, 25, and 53 | 67, 33, 17, 29 | 33, 34, 19, 17, 29, PAN, and unfertilized Sac4 |

### Expression profiling of seed-specific genes regulated by bZIPs

Gene investigator was used to analyze the transcript profiles of genes regulated by bZIP TFs under study here during different developmental stages of *Arabidopsis* (Supplementary figure S1). Expression data revealed the higher and continuous expression of bZIP53 throughout the developmental stages of plant starting from seedling to seed maturation. Similarly, expressions of bZIP10 and bZIP25 were also observed that overlaps with bZIP53 expression. The expression of target genes of bZIP53, bZIP10, and bZIP25 namely, *2S2*, *LEA76*, *ASN1*, *CRA1*, and *CRU* increased during seed development and maturation (Supplementary figure S1). However, expression of *ProDH*, a direct target gene of bZIP53 was found to be higher in the mid stage of development, which eventually decreased during the seed maturation (Weltmeier et al. 2006). bZIP TFs like bZIP39 and bZIP72, which do not form heterodimer with bZIP53 and its dimerizing partners are also involved in the seed development and maturation (Alonso et al. 2009; Belmonte et al. 2013; Bensmihen et al. 2005; Jain et al. 2018; Jain et al. 2017; Le et al. 2010). *bZIP72* has a profound expression in the cotyledon while *bZIP39* (ABI5) was found in the embryonic part of seed (Belmonte et al. 2013; Le et al. 2010). Although, *bZIP39* showed higher expression in the later stage of seed development and maturation but the target gene SHB-1 was downregulated in the corresponding phase (Cheng et al. 2014). This emphasizes that bZIP39 is also involved in other biological process and can heterodimerizes with other bZIP TFs and participate in seed development and maturation.

To gain an insight on the impact of A-ZIP53 over the expression of corresponding genes of target TFs including bZIP53, bZIP10, and bZIP25 (Lam et al. 2003; Weirauch et al. 2014; Weltmeier et al. 2006) transgenic *lines* were subjected to gene expression analysis. qRT-PCR was used to analyze the expression of seven target genes (*2S2, CRU, LEA76, ProDH, ASN1, CRA1,* and *HSD1*), and a non-target gene (*SHB-1*) (Alonso et al. 2009; Cheng et al. 2014; Weirauch et al. 2014). Leaves of the transgenic lines from the T-1 generation and immature siliques and seeds from T-2 generation were taken for the gene expression analysis (Figure 4A and 4B). The expression of *bZIP53* increased several folds in transgenic lines compared to the wild type (Figure 4). It could be to compensate the requirement of bZIP53 in transgenic, which is not available due to heterodimerzation with A-ZIP53. The expression of target genes of bZIP53, bZIP10, and bZIP25 including *CRU, ASN1, CRA,* and *HSD1* were downregulated in both generations (Figure 4A and 4B). The expression of seed storage albumin (*2S2*) and late embryogenesis accumulating 76 (*LEA76*) were not observed in the T-1 generation (Figure 4A) whereas both genes were downregulated in the T-2 generation (Figure 4B). The expression of *ProDH*, which is a direct target of bZIP53 is higher in the T-1 generation while expressed less in the T-2 generation (Figure 4A and Figure 4B) (Weltmeier et al. 2006). Thus, it could be inferred that, several fold higher expression of bZIP53 might overcome the inhibitory effect of the A-ZIP53 and led to the higher expression of the *ProDH*. Previously, we demonstrated the specificity of the A-ZIP53 and showed it does not heterodimerize with bZIP39 and bZIP72 in vitro (Jain P et al. 2017). The higher expression of the SHB-1, a direct target of bZIP39 confirmed the specificity of A-ZIP53. It shows the specificity and efficacy of DN protein to regulate the redundant behavior of target bZIPs.

**Figure4.**
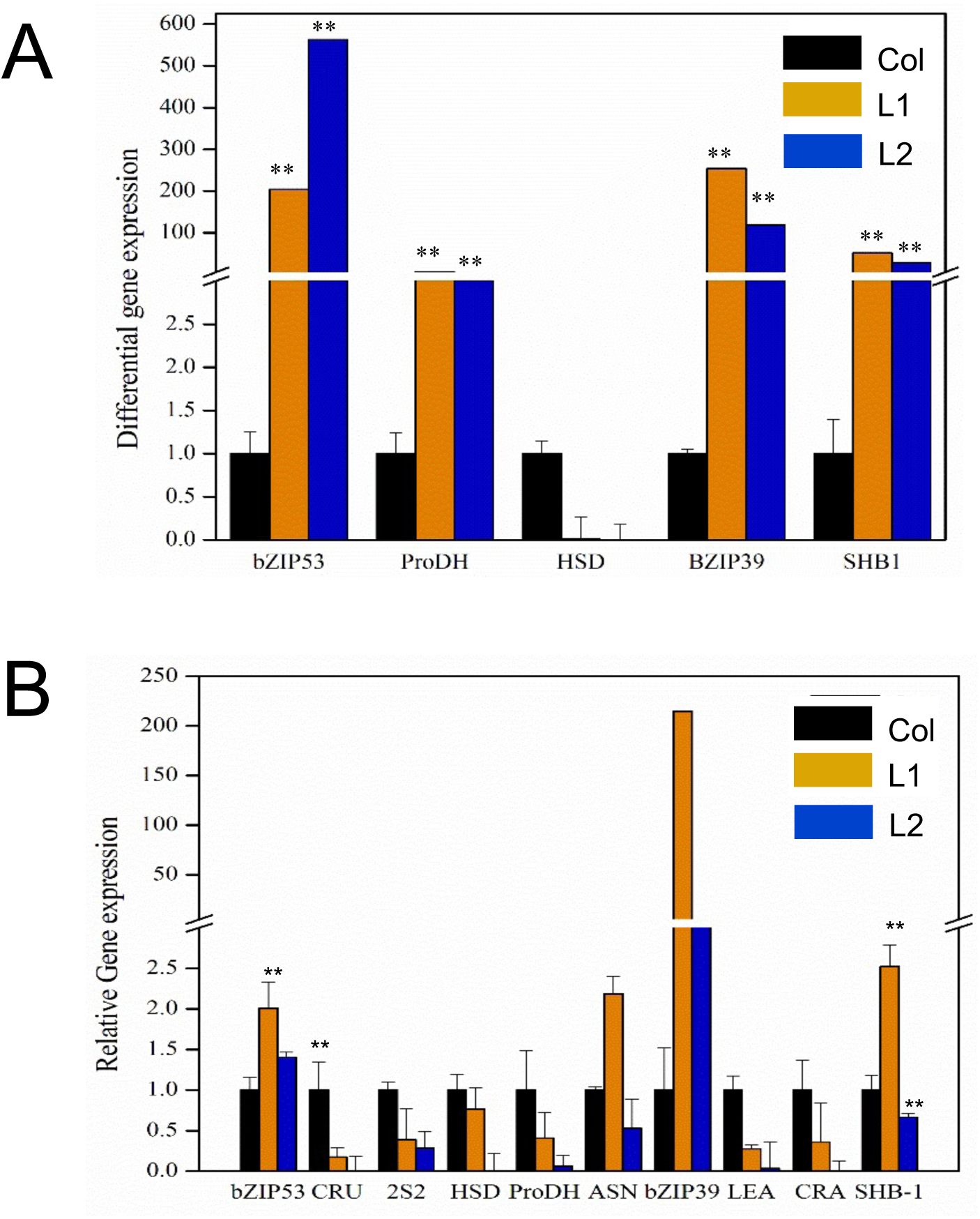
(A) qRT-PCR revealed the expression of bZIP53, bZIP39, target (2S2, LEA, CRU, ProDH, ASN1, CRA1, and HSD1) of *bZIP53* and non-target gene (SHB -1) from the leaves of four weeks old transgenic and immature siliques and seeds of six week old plants.

### Varied reproductive phase parameters of transgenics

To investigate the effect of A-ZIP53 on the reproductive phase of plants, A-ZIP53 expressing plants were studied together with the mutants of *bZIP53*, *bZIP10*, and *bZIP25* and wild type *Arabidopsis* (Supplementary Figure S2). *bZIP10* and *bZIP25* are reported to be involved in early stages of seed development (Lara et al. 2003) while the role of *bZIP53* is reported in the later stages of seed maturation (Alonso et al. 2009). A transgenic line, which has lower expression of A-ZIP53 was used for further analysis. Initially, this transgenic line has delayed growth like its predecessor and has lesser rosette diameter compared to mutants of *bZIP53*, *bZIP10*, *bZIP25*, and wild type (Supplementary Figure S2F) but later no significant differences were observed in plant height, growth, and leaves compared to mutants and wild type plants. Immature siliques and seeds of transgenic were subjected for qRT-PCR that showed the lower expression of genes involved in seed development and maturation (Supplementary Figure S2G). The expression of *bZIP53* was many fold higher compared to the wild type. A significant higher expression of the *HSD1* and *bZIP39* also observed in the A-ZIP53 expressing transgenic lines.

Transgenic plants were analyzed for phenotypic variation including developed flower, silique, and mature seed compared to mutants of the *bZIP53*, *bZIP10*, *bZIP25*, and wild type *Arabidopsis*. Transgenics has reduced flower size compared to the wild type and mutants of bZIPs while flower development was like wild type (Figure 5A). Significant differences between flower of transgenics and mutant of *bzip10* and *bzip25* were observed but no significant difference were seen compared to *bzip53* mutant (Figure 5). The diameter of transgenic flower was significantly less compared to wild type and mutants of *bzip10* and *bzip25* (Figure 5B). Furthermore, transgenic has smaller siliques compared to the wild type and mutants (Figure 6A and 6B) and number of siliques per 0.5 gm of weight is more compared to wild type (Figure 6C). Seeds of transgenic were small and shriveled (Figure 7A and Figure 7B). The seed weight of mutants and transgenics was less compared to the wild type (Supplementary figure S3). Small flower size, shorter silique length, and lesser number of seeds represents the impact of *A-ZIP53* on reproductive and seed development stage of the plant. Additionally, the length and width of seeds were significantly less compared to wild type *Arabidopsis* (Figure 7A and 7B). These results signify the effect of A-ZIP53 on the functioning of *bZIP53* and its heterodimerizing partners, which are involved in seed development and maturation.

**Figure 5.**
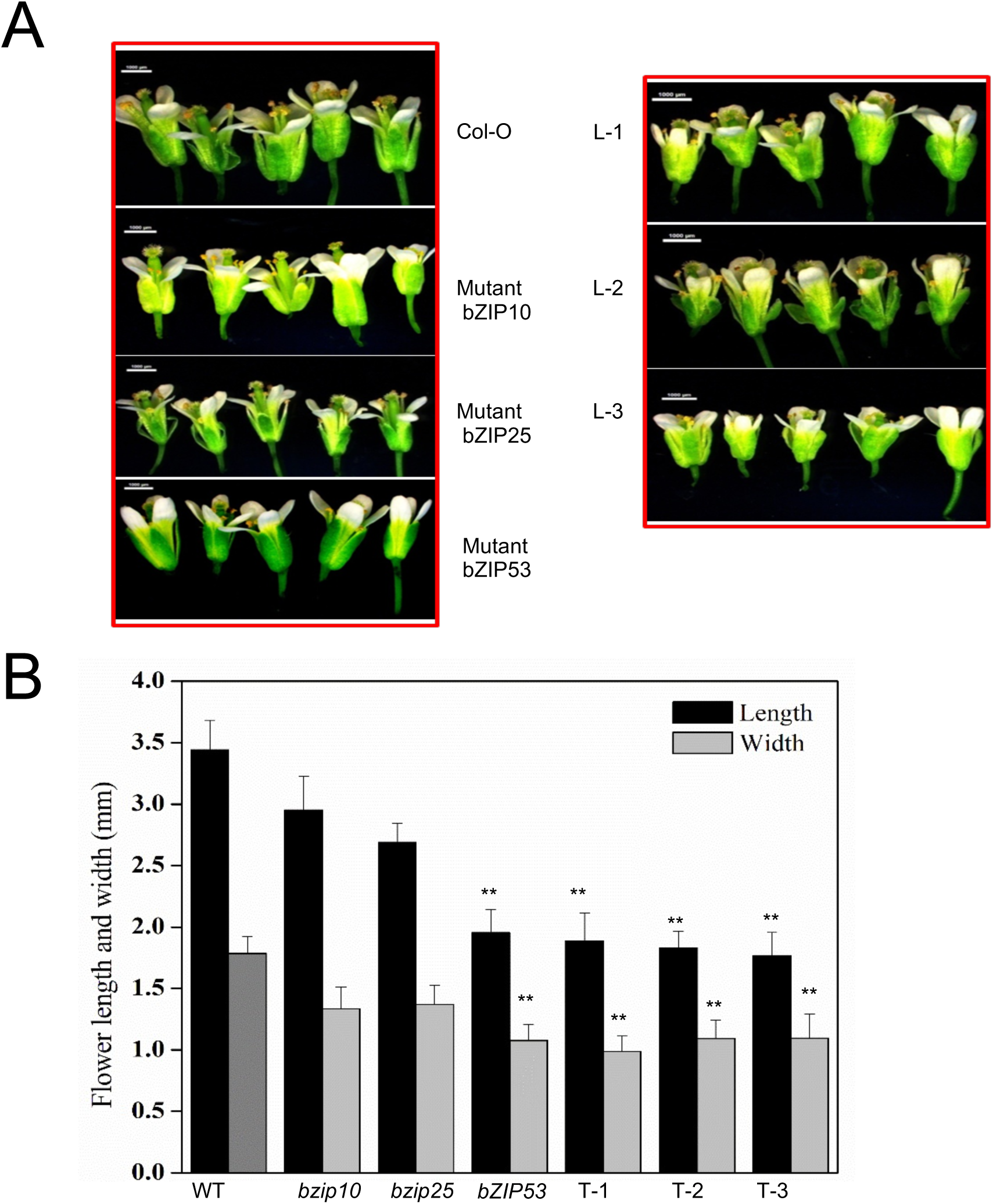
(A) Differences between length and size of flowers of mutants (*bzip10*, *bzip25*, and *bZIP53*) and transgenic of six weeks old plant **(B)** Significant differences between length and width of transgenic and mutants compared to wild-type.

**Figure6.**
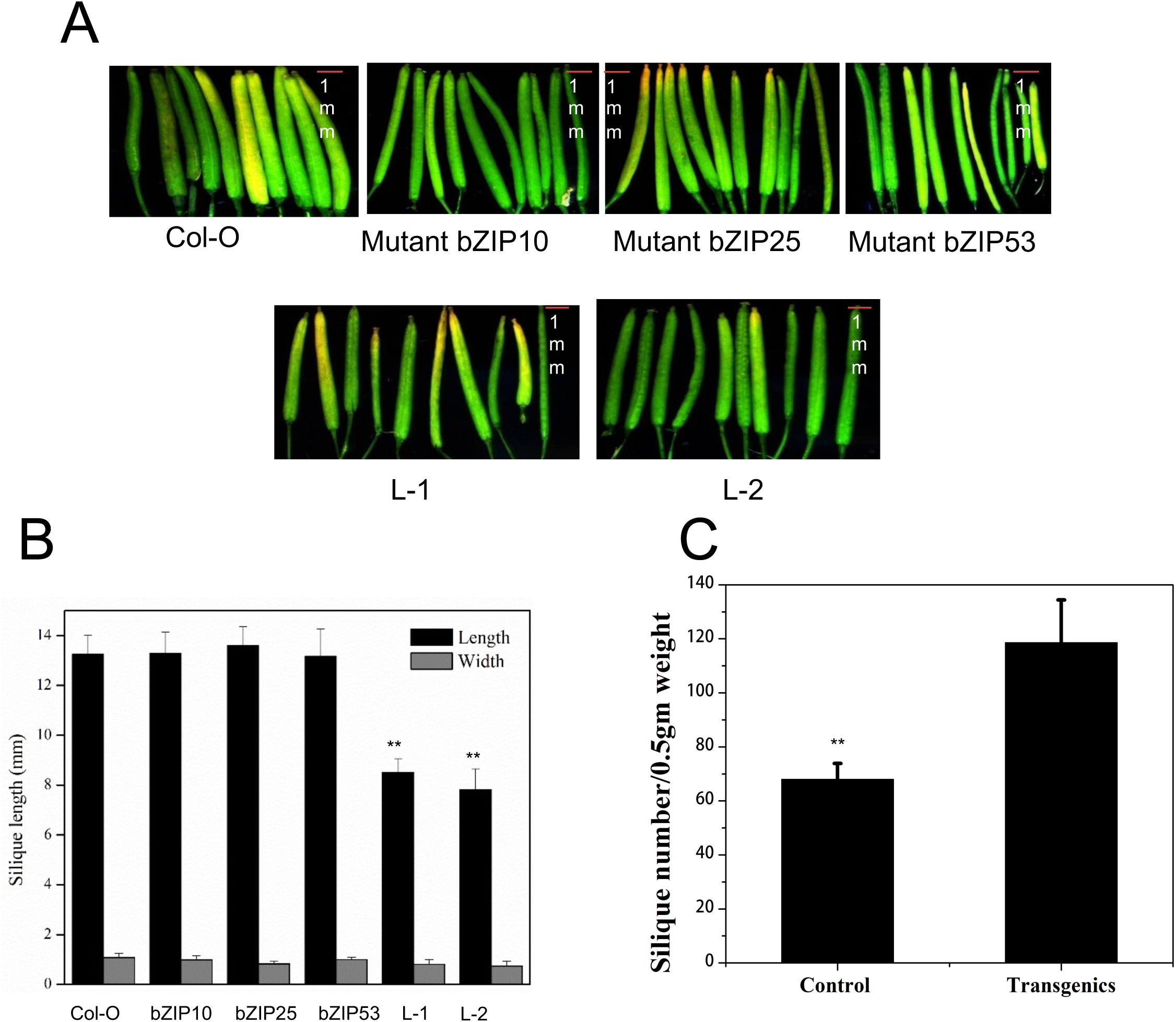
**(A)** Silique size of transgenic were smaller compared to mutants and wild type. **(B)** Length and width of siliques were compared. Error bar represent ±mean and SD of siliques (n = 8-12) **(C)** Number of siliques per 0.5 gm of silique weight were more in transgenics compared to wild type. (± mean and S.D. of three independent biological replicates).

**Figure7.**
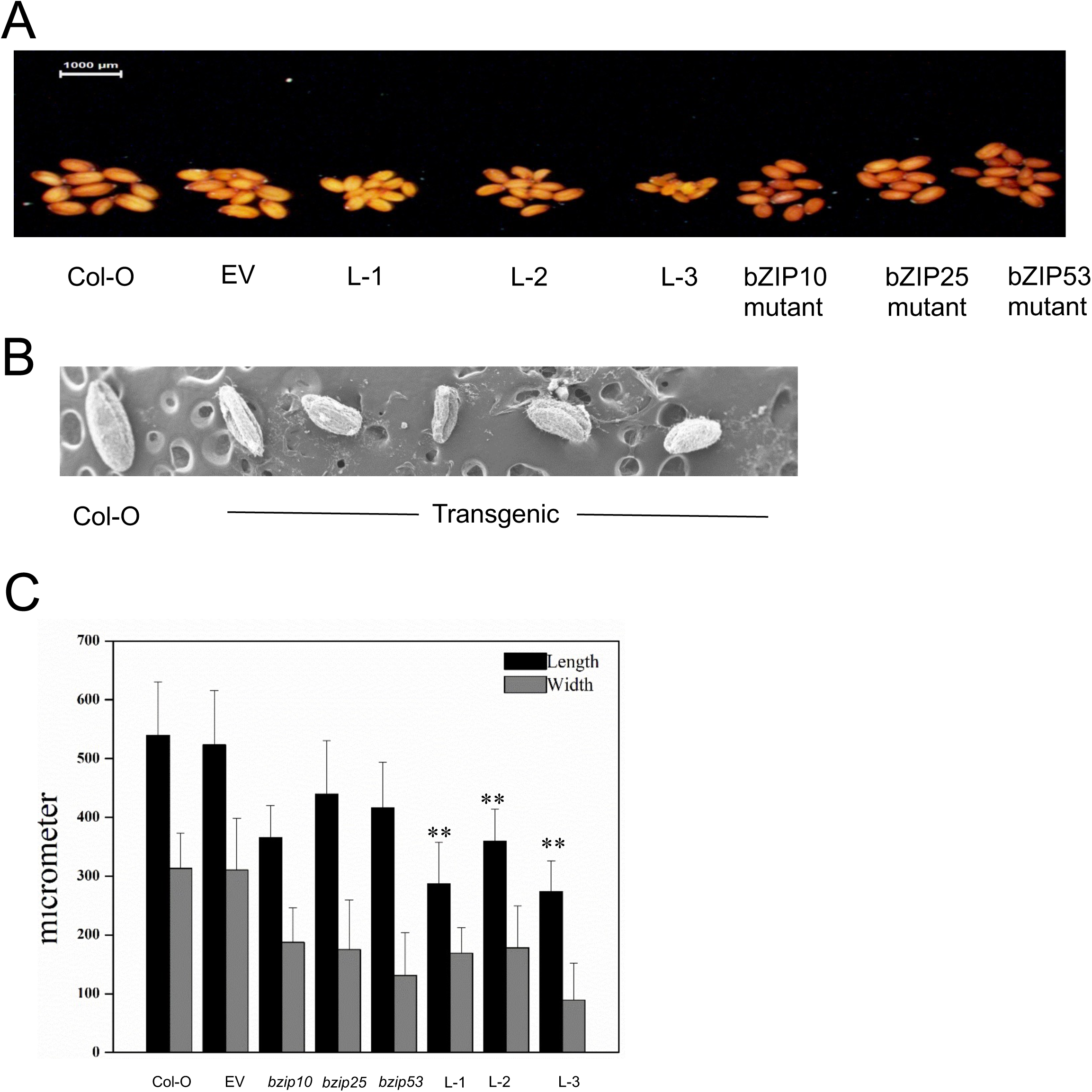
Differences in the seed size of transgenic, wild type, and mutants (*bZIP53*, *bzip10*, and *bzip25*) **(A)** Representative pictures of wild type, mutants and transgenic seeds **(B)** Seed of transgenic are smaller size compared to mutants and wild type (Error bar represent ±mean and SD of siliques (n = 25).

### A-ZIP53 restrict the expression of seed-specific genes

To understand the transcriptome dynamic in the A-ZIP53 expressing plants compared to the wild type, RNA-seq analysis was performed. High quality reads were mapped on reference genome of *Arabidopsis thaliana*, which ranged from 90 % to 92 %. Two biological replicates were observed to understand the effect of A-ZIP53. The transcriptome data revealed 1029 differentially expressed genes (DEG) in which 71.62 % (737 out of 1029) were upregulated compared to 28.38 % (292 out 0f 1029) were downregulated genes (Figure 8) in A-ZIP53 expressing transgenic plants.

**Figure 8.**
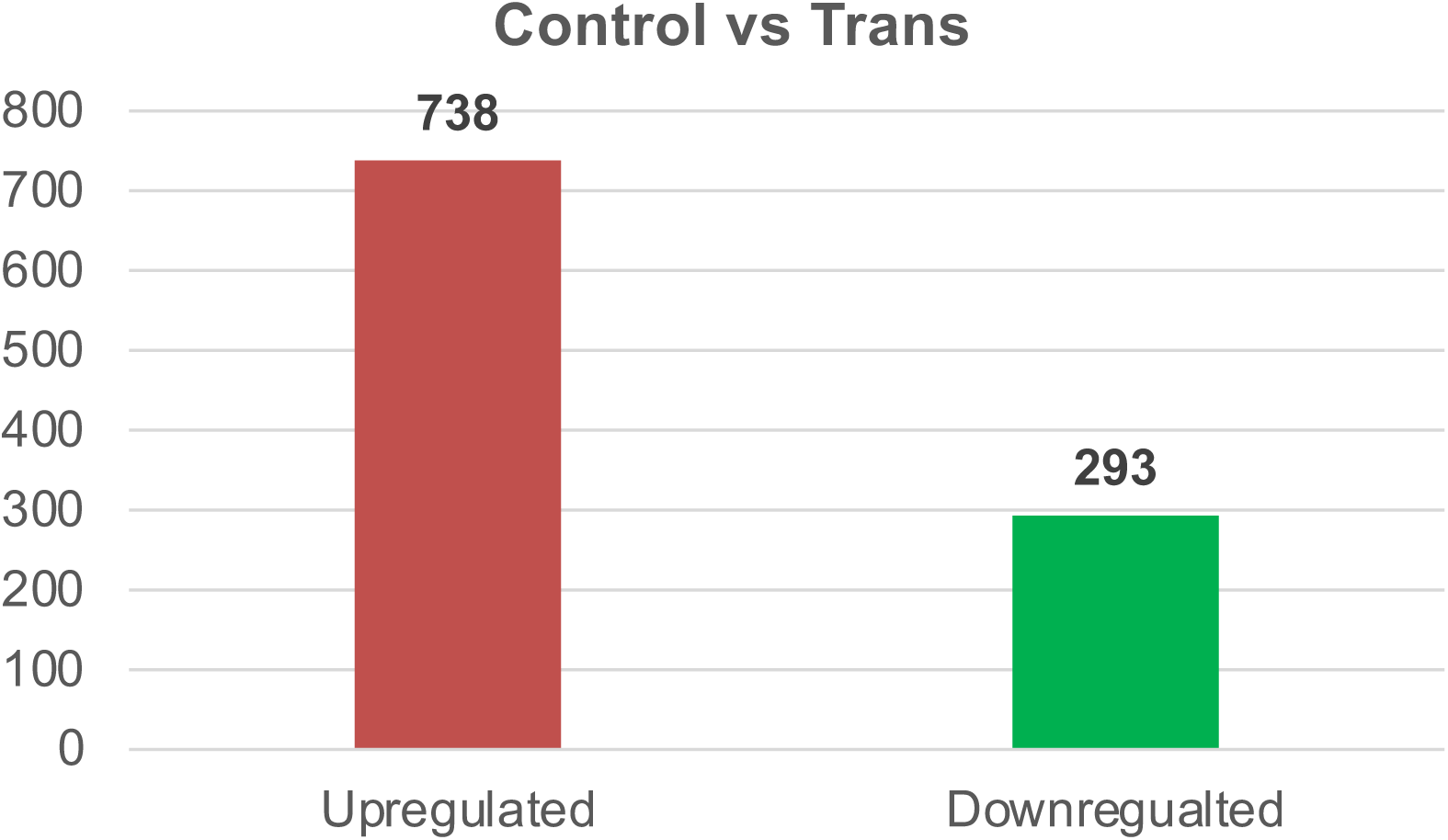
Differential expression of genes (DEG) based on analysis from two independent biological replicates of wild type and A-ZIP53 expressing transgenic.

The assembled data was functionally categorized using the agriGO gene ontology (GO) tool (Figure 7). 20,764 unique transcripts, with the FDR of 0.05 were examined with the GO tool. The knocking down of *bZIP53* and its heterodimerizing partners have a profound effect on the expression of corresponding genes. Down regulated transcripts were categorized into GO terms that participate into biological, cellular, and molecular functions (Supplmentary Table S4). Most of the downregulated GO terms were identified to be related to genes, which are involved in gamete formation, seed development, seed maturation, seed storage protein synthesis, reproduction, and other biological processes. 48.8% genes in the biological, 20.8% in the cellular, and 32.2% genes involved in the molecular functions have been identified (Figure 9). Genes related to developmental process, hormone metabolism, and DNA binding transcription factor s activities were down regulated (Figure 9), which suggests the A-ZIP53 role inthe development related pathway.

**Figure 9.**
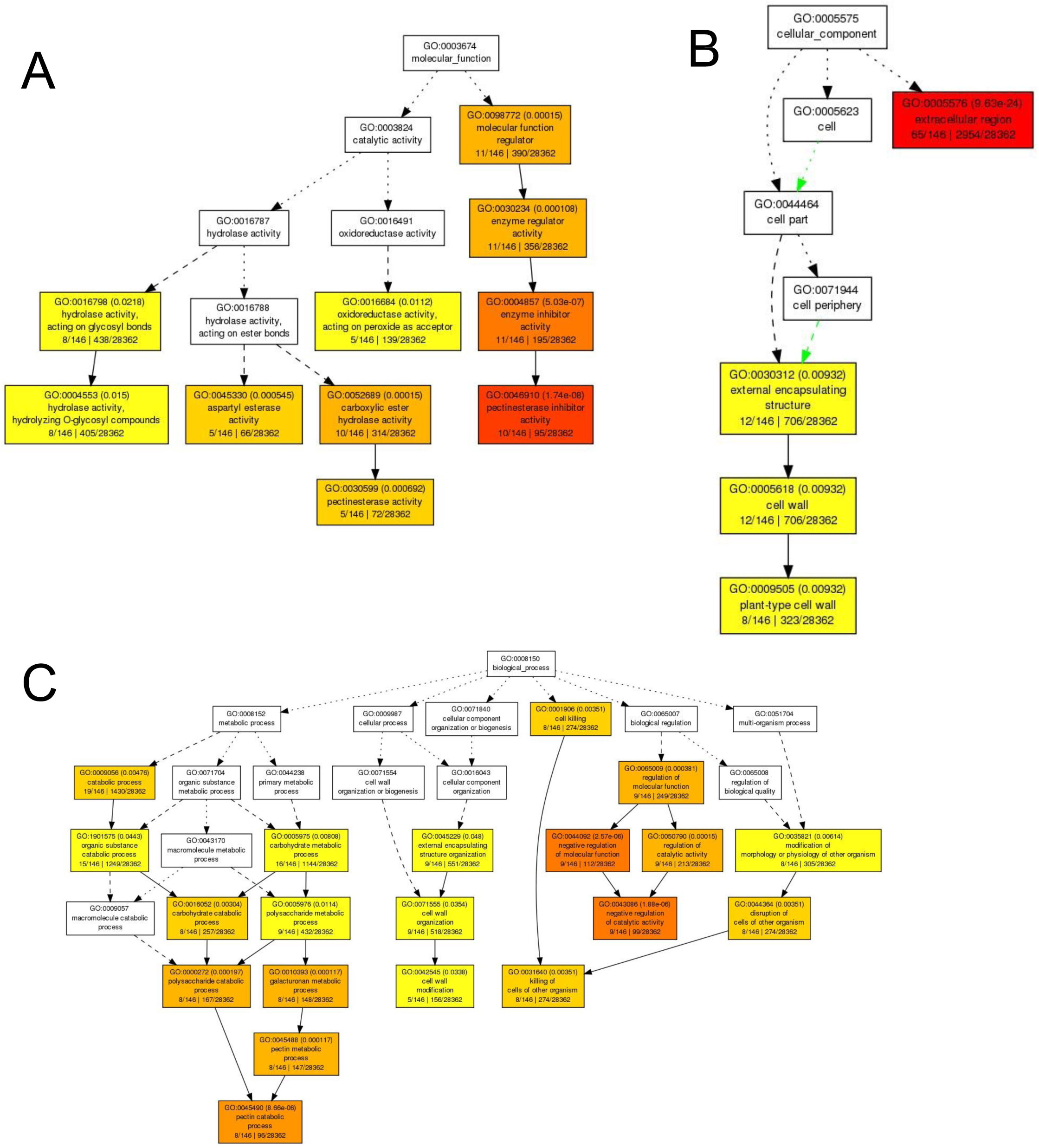
Gene Ontology enrichment analysis (for downregulated genes w.r.t. upregulated genes) resulted in selection of large number of enriched classification terms. Similar in case of enriched GO terms in upregulated genes w.r.t. all genes in Arabidopsis. **A)** Genes involved in molecular function. **B)** Genes involved in cellular function. **C)** Genes involved in biological function

The lower expression of genes related to gamete and seed development pathway including Late embryogenesis abundant protein (LEA) family, ECA1 gametogenesis related family protein, maternally expressed family protein, seed storage 2S albumin superfamily protein and others substantiate the A-ZIP53 effect on the target genes. Highlighting the redundant behaviour of TFs, genes involved in stress including salt stress are also found to be differentially expressed in A-ZIP53 expressing transgenic plants.

### Decoding of target genes and regulatory network

Data generated from this study has helped to identify the putative target genes of *bZIP53* and its dimeric partners involved in regulating seed storage protein, gamete development, transcription factors, and both biotic and abiotic stresses.

DN protein A-ZIP53 has served to capture different proteins, which might act synergistically in different biological process like correlation between hormone signaling and TFs binding. One such example is bHLH-MYB complex in jasmonic acid -mediated stamen development and seed production (Qi et al. 2015). Transcriptome data revealed the higher expression of bHLH and MYB genes, which might be to balance the hinderance in function of other TFs including bZIPs. A-ZIP53 expressing transgenics have a retarded growth and small flower and seed phenotype. It might be due to the DN interferes with function of other regulating TFs like AGL100 (Bemer et al. 2010), MYB24 (Qi et al. 2015), ARF4 (Liu et al. 2018), and bZIP53 (Alonso et al. 2009) in an indirect manner, which might be working synergistically in flower development, gamete formation, and seed development. Disturbing the action of one TFs eventually imbalances the whole complex transcriptional machinery. Decipher the complex gene regulatory machinery governed by other unknown factors through DN protein is helpful that not be possible using other loss of function mutation techniques.

The analysis of transcriptome revealed the complex combinatorial network of TFs including bZIPs (bZIP1 and bZIP53), MYB (MYB2, MYB19, MYB24, MYB27, MYB35, MYB51, and MYB59), MADS (AGL100), ARF (ARF4), and WRKY (WRKY6, WRKY15, WRKY28, WRKY33, WRKY47, WRKY48, WRKY66, and WRKY75), which might be working in a combinatorial fashion in stress and seed development.

### Possible targets of A-ZIP53 during seed maturation

Previous study by Alonso et al., 2009 showed that bZIP53 is a key regulator for seed maturation that can form heterotypic interaction with bZIP10 and bZIP25 (Alonso et al. 2009). However other bZIPs are also reported to be involved in seed development and maturation, including bZIP39 (Bensmihen et al. 2005; Cheng et al. 2014; Dekkers et al. 2016). bZIP39 is also involved in floral transition (Wang et al. 2013), which signifies the functional redundancy like bZIP53 (Alonso et al. 2009; Dietrich et al. 2011; Hartmann et al. 2015). Functional redundancy of bZIP depends on the different dimerizing partner selection.

Our finding showed that A-ZIP53 can form the heterotypic interaction with the bZIP53, bZIP10, and bZIP25 *in vitro* and *in vivo* (Jain et al. 2018; Jain et al. 2017). In order to know other heterodimerizing partners of A-ZIP53, whole protein extract from immature siliques, immature seeds, and leaves were subjected to the immunoprecipitation followed by the mass-spectrometry (IP-nano LC-MS/MS (Material and Methods). bZIP proteins that were identified in more than one sample with at least one proteotypic peptide were considered as a high confidence candidate. Eight bZIP TFs (bZIP14, bZIP17, bZIP19, bZIP23, bZIP29, bZIP33, bZIP34, and bZIP69) were found in the study, which could be the interacting partners of the bZIP53 or A-ZIP53 (Table I). The similarity between the bZIPs (bZIP29, bZIP33, and bZIP67) precipitated from both samples confirm the efficacy and effectivity of A-ZIP53. In addition, to confirm the dimeric specificity of A-ZIP53 with target bZIPs, the total protein soup of wild-type Arabidopsis was incubated with pure protein A-ZIP53 followed by IP-MS. The annotated peptides were related to bZIP33, bZIP29, and bZIP53, which confirms the precipitation of similar bZIPs in all samples and efficacy of A-ZIP53. Amino acid sequences of immunoprecipitated bZIPs and their propensities to form dimeric interaction with bZIP53/A-ZIP53 also analyzed (Supplementary figure S5, S6, and S7). Charged amino acids at g and e’ positions in the leucine zipper forms an interhelical g ↔ e’ (Jain et al. 2017). The parameter was used to quantify the putative interactions between immunoprecipiated bZIPs with bZIP53 and A-ZIP53. Immunoprecipitated bZIPs have more putative attractive heterodimeric interactions with bZIP53 and A-ZIP53 except bZIP29 and bZIP33, which are common for all samples and more repulsive interhelical g ↔ e’ interactions (4 for each) (Supplementary Figure S6B and S7B). It might be that these are the part of a novel interacting pathway involved in different developmental process.

### A-ZIP53 inhibits the DNA binding of target bZIPs in the transient transfection using Arabidopsis protoplast

Transient transfection with the Arabidopsis protoplast was used to probe the affectivity of A-ZIP53 against target bZIPs identified by the IP-MS. For transfections, construct of 2S2 promoter which regulates the expression of GUS was co transformed with effector plasmids (*A-ZIP53*, and other bZIPs (*bZIP14, bZIP17, bZIP19, bZIP29, bZIP34, bZIP53,* and *bZIP69*) and control plasmid (NAN), which were under the control of 35S promoter (Alonso et al. 2009; Rishi et al. 2004; Weltmeier et al. 2009). A higher GUS/NAN activity was observed in the co-transfections of when co-transformed with the bZIPs plasmids (bZIP53|bZIP69). It shows a positive heterodimeric interaction between bZIP53 and bZIP69 in the presence of DNA. However, A-ZIP53, which is a DN of bZIP53 heterodimerizes and inhibit the DNA binding of target bZIPs, which results in reduced GUS/NAN activity, which shows the other mode of interaction between immunoprecipitated bZIPs and A-ZIP53 (Figure 10).

**Figure10.**
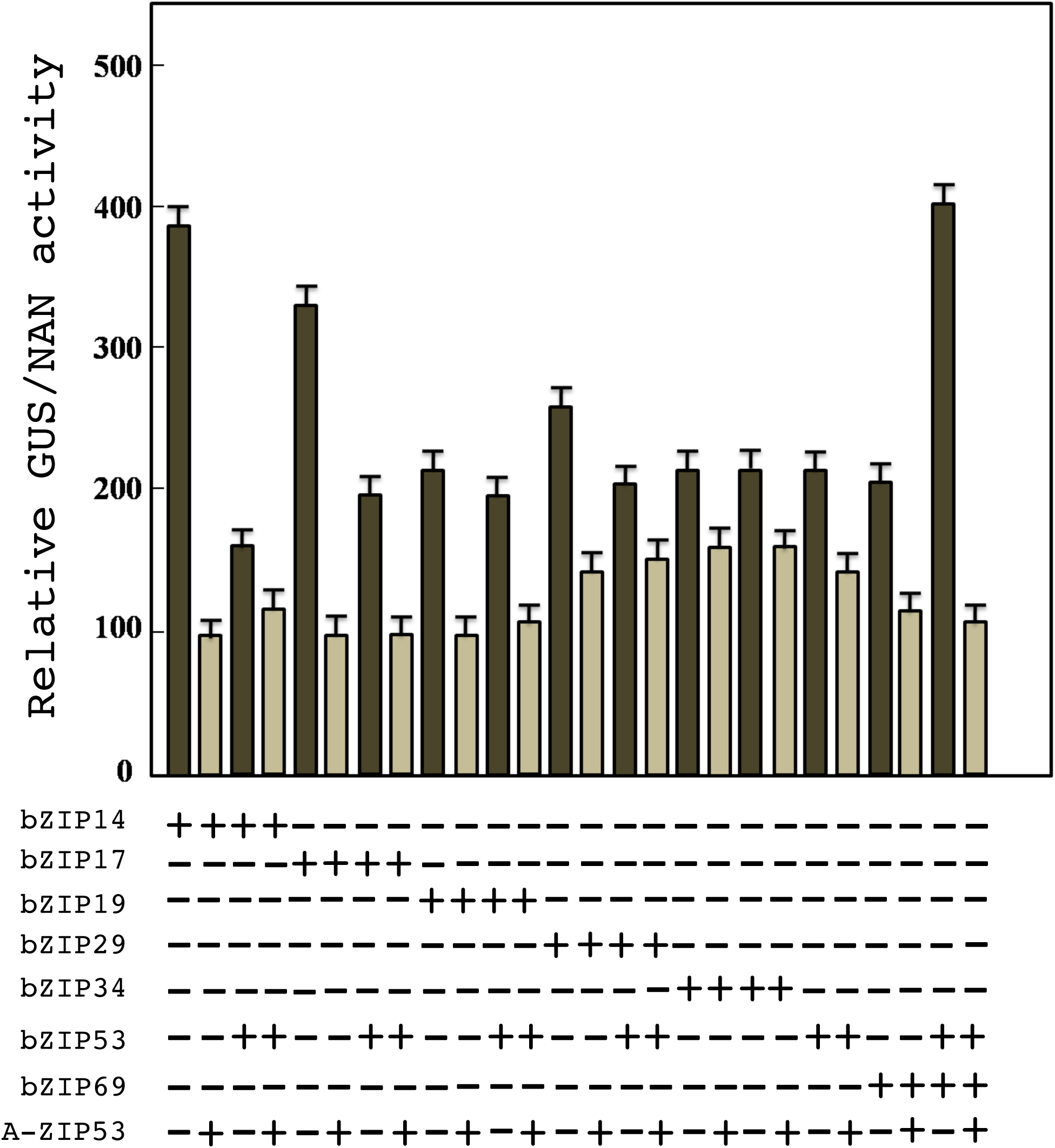
A-ZIP53 hinders the DNA binding of the bZIP14, bZIP17, bZIP19, bZIP29, bZIP34, bZIP53, and bZIP69 and their heterodimer with the bZIP53 mediated reporter gene activity in transient trsnsfection assay using Arabidopsis protoplasts.(**A**) Plasmid coding for bZIP53 can transactivate the *GUS* reporter gene under the control of the 2S2 promoter that contains a G-box binding site. ** represent P < 0.01. Protoplasts were co-transfected with plasmids coding for bZIPs and A-ZIP53. Transient expressions of A-ZIP53 inhibited the bZIPs-mediated *GUS* reporter activity in dose-dependent manner. Reporter activity was inhibited in the presence of 3 molar excess of A-ZIP53 plasmid suggesting that A-ZIP53 can compete with G-box for bZIPs binding. Error bars represent standard deviation of three independent experiments.

## Discussion

Various studies in the past have been done to decipher the molecular network for seed maturation however, less attention has been paid to regulate the redundant behavior of transcription factors. In this study, we have established the molecular worthiness of a novel designed dominant negative protein A-ZIP53 to regulate the dimerization of target bZIP transcription factors. A-ZIP53 is previously reported to form heterotypic interaction with target bZIP TFs i.e., bZIP53, bZIP10, and bZIP25, which are involved in seed maturation (Alonso et al. 2009; Jain et al. 2017). The overexpressed A-ZIP53 specifically form a heterotypic interaction with target bZIP proteins and hinder their DNA binding activity. The p35S:A-ZIP53 plants have retarded growth and produces unviable seeds, indicating the efficacy of designed protein against target bZIP TFs.

Earlier, we have functionally validated A-ZIP53 and its derivatives against target bZIP TFs in *Arabidopsis* (Jain et al. 2018; Jain et al. 2017). A-ZIP53 has a dimerization domain of bZIP53 and a designed polyglutamic rich acidic extension at its N-terminal. The design strategy has been adapted from the idea that the acidic extension mimics DNA and provide an alternate binding site for target bZIP TFs (Krylov et al. 1995). This acidic extension stretches the leucine zipper extension to A-ZIP53|bZIP heterodimer. The functional significance was examined by constitutive expression of *A-ZIP53* under control of CaMV35S promoter. Arabidopsis plants expressing a novel A-ZIP53 partially mimics the phenotype of bZIP53 overexpressing lines, with a dwarf and delayed bolting phenotype (Alonso et al. 2009). The differential expression of A-ZIP53 confirms the efficacy and potent effect on plants (Figure 1). A-ZIP53 has a dimerization domain of bZIP53 that belongs to class S1 bZIP TF (Jakoby et al. 2002). Earlier reports confirmed the putative heterotypic interaction between bZIPs of group S1 and group C namely bZIP9, bZIP10, bZIP25, bZIP63, and others using yeast two hybrid and *in vitro* DNA binding assays, which have a prominent role in growth and development (Alonso et al. 2009; Dietrich et al. 2011; Dröge-Laser et al. 2018; Ehlert et al. 2006; Hartmann et al. 2015; Kang et al. 2010; Weltmeier et al. 2006). The network of class C/S1 bZIPs is less disordered and have heptads with a high helical tendency that favors their dimerization (Deppmann et al. 2006; Jakoby et al. 2002). Further, heterodimers of class C/S1 bZIPs have lesser stabilizing forces resulting in the weaker stability (Llorca et al. 2014). It prompted us to imply the DN protein A-ZIP53 that efficiently and stably forms a heterotypic interaction with the class C/S1 bZIP TFs. The designed acidic extension prolongs the dimerization interface into the DNA binding region and provides two magnitude higher stability to the A-ZIP53|bZIP heterodimer complex (Jain et al. 2017). Excess of A-ZIP53 extends its specific dimerization and tendency to target bZIPs, which is less abundant. It makes A-ZIP53 an effective competitor in a stoichiometric environment.

It was shown previously that the O2, bZIP10, and bZIP25 related to the group C bZIPs interacts with ABI3 during seed maturation and bZIP53 enhanced the activation of the heterodimer complex in transient transfection assay (Alonso et al. 2009; Ehlert et al. 2006; Schmidt et al. 1990; Weltmeier et al. 2009). The specificity of A-ZIP53 against class C/S1 bZIPs was validated by transient transfections, which showed the efficacy of DNs to overcome biological redundancy (Satoh et al. 2004; Weltmeier et al. 2006) and stresses (Dietrich et al. 2011; Hartmann et al. 2015).

Redundant behavior of bZIP TFs is due to their interacting partners and A-ZIP53 has an edge to decipher them. It is reported that bZIPs has strong transactivation properties and the heterotypic interaction with the promiscuous DN like A-ZIP53 may deter their potential binding to cognate DNA binding sites. These DNA binding sites can play an active role for the cooperative interactions between bZIP TFs (Jain et al. 2017; Jolma et al. 2013). These can be varied for the individual TFs and it may be due to the different heterodimerizing partners like the bZIP53 TF, which has affinity to the G-box (AC<u>ACGTG</u>AT) and the C-box (CC<u>ACGTC</u>GC) as shown by the Protein-DNA binding assays and the DAP-seq (Supplementary Table S1) (Alonso et al. 2009; Ezcurra et al. 2000; Jain et al. 2017; O’Malley et al. 2016; Pedrotti et al. 2018). DNA mimicking acidic extension of A-ZIP53 provides an alternate binding site for bZIP TFs and hinders their interaction with partner bZIPs and DNA (Jain et al. 2017). A-ZIP53 has more tendency to interact specifically with target bZIP proteins compared to DNA (4.6 kcal mol^-1^ dimer^-1^) (Jain et al. 2017). It deters the function of target bZIP TFs and regulates their redundant behavior by forming heterodimer like the *bZIP53*, which is a direct target of *A-ZIP53* (Alonso et al. 2009; Hartmann et al. 2015; Jain et al. 2018; Jain et al. 2017). *bZIP53* expression is not restricted to seeds and can be observed in vegetative tissues (Supplementary figure S1). Arresting its heterodimerization results in the abnormal phenotype like retarded growth and shriveled seeds (Figure 2B and Figure 2C)(Alonso et al. 2009; Dietrich et al. 2011; Hartmann et al. 2015).

Expression of genes including *ProDH*, *ASN1* are reported to be directly regulated by *bZIP53* (Baena-González et al. 2007; Hanson et al. 2008; Satoh et al. 2004; Weltmeier et al. 2006). ProDH is involved in hypo-osmolarity response and ASN1 is known to enhance seed storage protein (SSP) content and nitrogen status of food (Lam et al. 2003; Satoh et al. 2004; Weltmeier et al. 2006).Studies done on bZIP53 overexpressing transgenic lines have revealed higher expression of *MAT* and *LEA* genes like *2S2*, *CRU3*, *CRA1*, *HSD1*, *LEA76*, *ProDH*, and the *ASN1* (Alonso et al. 2009; Baena-González et al. 2007; Lam et al. 2003; Satoh et al. 2004; Weltmeier et al. 2006). Significantly lesser expression of *MAT* and *LEA* genes was observed in the immature siliques and seed of the T-1 and T-2 generation of the A-ZIP53 expressing transgenic lines, which could be a reason for the abnormal and shriveled seeds phenotype (Figure 7A and 7B) (Tunnacliffe and Wise 2007). A-ZIP53 regulates the C/S1 network and deter their interaction with the cognate DNA binding sites. A-ZIP53 is highly specific to its target as no such negative effect was observed on the expression of bZIP39 and its target gene *SHB-1* (Cheng et al. 2014; Jain et al. 2017). The higher expression of *bZIP53* was observed in the transgenic lines of the T-1 and T-2 generation, which could be to compensate the unavailability of the *bZIP53* to other dimeric partners. The effects of reproductive parameters were subjected for the analysis..

It was observed that A-ZIP53 expressing transgenic have significantly smaller flower, shorter siliques, and shriveled seeds compared to wild type (p <0.01), mutants of *bZIP53*, *bZIP10*, and *bZIP25* (Figure 5, Figure 6, and Figure 7). Phenotypic examination of the A-ZIP53 expressing T-3 transgenic lines have similar growth as wild type and mutants but have lower expression of *MAT* and *LEA* genes. In seed maturation, ABI3 is an important regulator that interacts with the bZIP10 and bZIP25. A ternary complex was reported between the ABI3, bZIP10, bZIP25, and bZIP53 is a key regulator in seed development and maturation (Alonso et al. 2009; Lara et al. 2003). Expression of the A-ZIP53 could restrict the DNA binding of ternary complex and restrict the function of non-target protein.

Transcriptome analysis of A-ZIP53 expressing transgenic lines revealed the down regulation of multiple genes including the LEA (log2FC -7.45) and 2S2 (**-**4.49), that are known to be crucial for the seed development and maturation (Supplementary table 8). These results signify the efficacy of A-ZIP53 against target bZIPs regulating genes involved in seed development and maturation. Additionally, number of crucial genes have been identified in this study, which might be involved in the growth, development or seed maturation pathway targets to uncover the complex cascade of seed development and maturation.

Through systematic transcriptomic analysis, a broad map of genes effected by A-ZIP53 has been discussed, which will help to uncover the complex cascade of seed development and maturation. These genes implicate in diverse processes including biotic (At1G19610) (Ascencio-Ibánez et al. 2008), and abiotic (At5g12030) (Lee and Seo 2019), DNA binding transcription factor (MYB24) (Qi et al. 2015), AGL100 (Bemer et al. 2010), WRKY33 (Wang et al. 2020) (and others), hormone signaling (Auxin-responsive GH3 family protein (Zheng et al. 2016), seed development and maturation (Late embryogenesis abundant protein (LEA) family protein (Candat et al. 2014).

Given the other interacting partners of bZIP TFs, a varied expression of cysteine rich peptides (CRPs) has been also observed. CRPs are reported in various biological processes, including mammalian and plant defenses, stress response, development and reproduction, and cell–cell communication (Marshall et al. 2011; Ostrowski and Kowalczyk 2015). They can form dimeric interactions (homo- or hetero-) and regulate diverse biological processes. The higher expression of CRPs in A-ZIP53 expressing plant signifies the relation between higher expression of bZIP protein and CRPs. A speculation can be made about a putative interaction between CRPs and bZIPs under salt stress as CRP (Xu et al. 2018) and bZIP1 or bZIP53 (Hartmann et al. 2015) both have a potential role in salt mediated stress. The lower expression of CRP (*AtPCP-Bδ-At2g16505*) was also observed, which regulates the pollen tube growth. It signifies the A-ZIP53 hinders the seed development pathway and have an effect on the post pollination events (Wang et al. 2017). The differential effect on the expression of CRPs, which might be a potential target of A-ZIP53 and eventually a possible heterodimeric partner of bZIP53 and other bZIPs has open a new-horizons about the molecular mechanism governed by bZIP TFs.

Immunoprecipitation followed by the mass spectrometry has confirmed the heterotypic interaction of A-ZIP53 with target bZIPs and its dimeric interacting partners (Table I). The precipitated bZIPs could be targeted in future to decipher the complex C/S1 or other novel cross network like B/I network (bZIP 17) (Jakoby et al. 2002). The transient transfection studies using protoplast confirmed the *in vivo* dimeric interaction between A-ZIP53 and target bZIPs (Figure 10). These results validate the DNA binding of bZIPs as a potential molecular target for functional regulation.

Based on findings from this study, a working model suggesting regulation of DNA binding and heterotypic interactions of target bZIP TFs is proposed (Figure 11). The function of bZIP53 in seed maturation depends on its heterodimeric partners bZIP10 and bZIP25. Ectopic expression of the bZIP53 leads to higher expression of genes involved in seed development and maturation. Designed novel DN protein A-ZIP53 could be used to restrict the DNA binding and heterotypic interaction of target bZIPs and inhibit their function.

**Figure 11.**
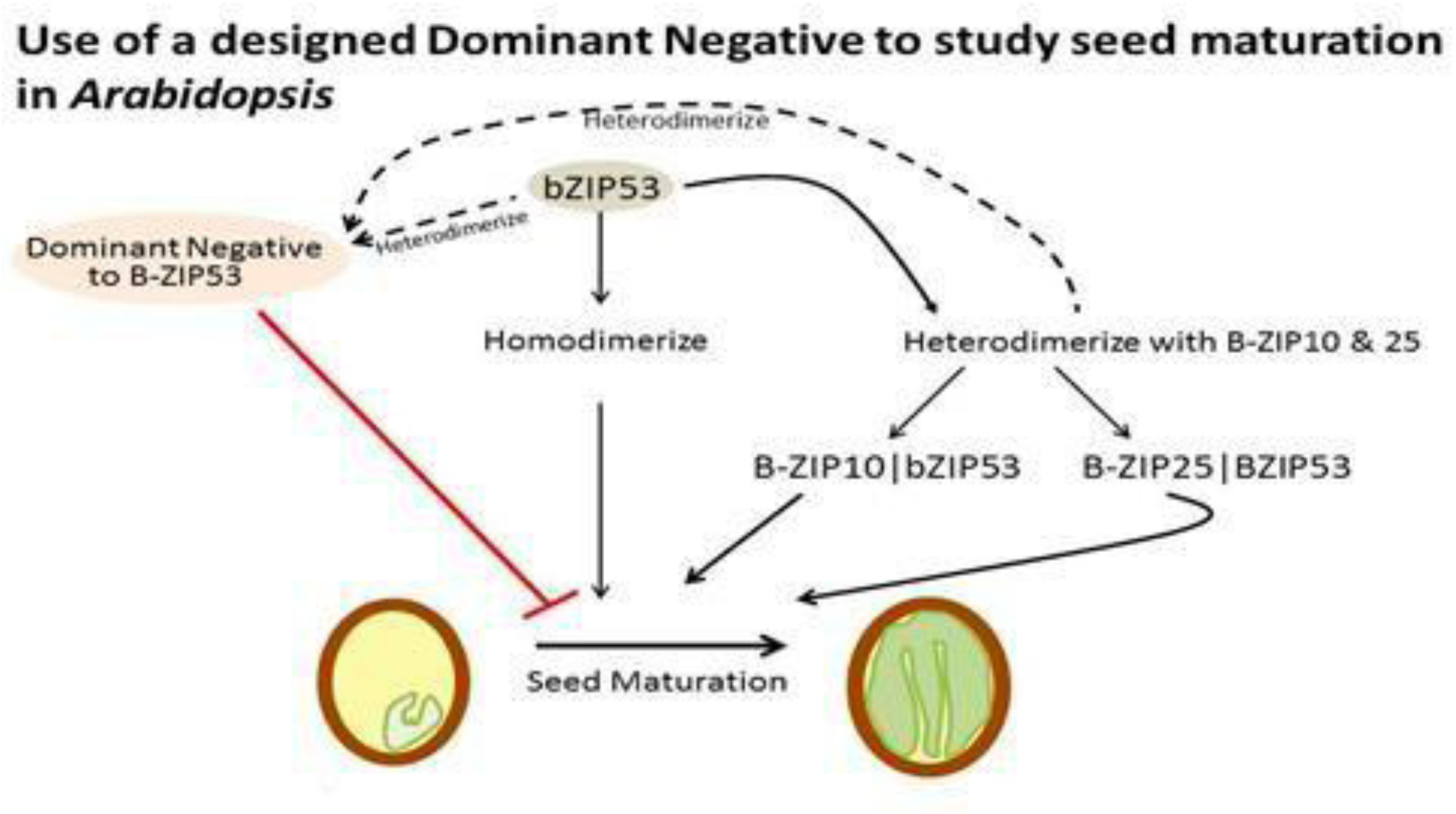
Model for heterodimeric regulation of *bZIP53*, *bZIP10*, and *bZIP25* involved in the seed maturation by the designed dominant negative protein *A-ZIP53*.

Moreover, to our knowledge this is the first time the efficacy of the *A-ZIP53* is reported to understand and regulate the redundant behviour of bZIP TFs in plants. While there are numerous techniques for loss of function studies like CRISPR-cas9 or RNAi, but their potential is limited to understand the redundancy. We therefore believe that the application of dominant negative proteins like A-ZIP53 could play a substantial role to understand the redundancy and unzip the complex cascade of signaling network. These interacting network of downstream signaling pathway could be a subject for further studies.

## Material and Methods

### Plant Material

Col -0 accession of *Arabidopsis thaliana* was used as a wild type in the present study. Seeds were surface sterilized and kept for stratification in dark at 4 °C for 2-3 days on half strength MS-agar (Millipore Sigma) plates. Seeds of mutants of *bzip10*, *bzip25*, and *bZIP53* were obtained from TAIR and germinated on the half strength MS agar plate. For germination, seeds were transferred in growth chamber under controlled condition of 16-h-light/8-h-dark photoperiod cycle, 22°C temperature,150 to 180 µmol m^-2^ s^-1^ light intensity, and 60% relative humidity. Three weeks old plants were used for the measurement of rosette diameter, and six weeks old mature plant were used for flower and silique size.

### Construct preparation and Plant Transformation

A-ZIP53 with the T-7 tag was amplified using the corresponding template and cloned as NdeI – EcoRI double digested fragment under the CaMV35S promoter in the pRI101AN vector.

To generate the *A-ZIP53* expressing trangenic lines, *Arabidopsis* (Col-0) plants were transformed with the floral dip method using *Agrobacterium* t*umefaciens* strain GV3101. Dipped plants were grown in a growth chamber under the standard growth conditions (22 °C under 16-h light/8-h dark photoperiod (150–180 μmol m^−2^ s^−2^). T1 transformed plants were selected on the 50 μg/ml kanamycin containing MS agar plate with 1 % sucrose for 7-10 days. Selected lines screened for the next generation and subjected for Western blotting.

### Protoplast isolation and transient transfection assay using BY-2 tobacco cell line and *Arabidopsis*

#### Protoplast isolation from BY-2 tobacco cell line

For protoplast isolation, BY-2 cell line suspension was centrifuged at 100 g for 5 minutes and 15 ml of packed cell volume was resuspended into 50 ml of protoplast isolation solution (7.4 gm/L CaCl_2_·2H_2_O, 1 gm/L NaOAc (anhydrous), and 45 gm/L mannitol supplemented with 1.2 % cellulose R10 (Millipore Sigma) and 0.6 % Macerozyme (Millipore Sigma), pH 5.7, filter sterilized with 0.22 μm filter). Suspension culture was transferred into three petriplate and incubated in dark with gentle shaking (100 rpm) at room temperature for 3-4 hrs. Protoplasts were centrifuged at 250Xg for 5 minutes, collected, and washed twice with protoplast isolation solution and centrifuged at 100 rpm for 1 minute. Pellet was resuspended in 10 ml of floating solution (99 mg/L myo-inositol, 2.88 gm/L L-proline, 100 mg/L enzymatic casein hydrolysate, 102.6 gm/L sucrose, 97.6 mg/L MES buffer, 4.4 gm/L MS salts, 1 mg/L thiamine-HCl, 370 mg/L KH_2_PO_4_, pH 5.7) and centrifuged at 250Xg for 10 minutes. Isolated protoplast on the top of solution were transferred to a new tube and 10 ml of W5 solution (154 mM NaCl, 125 mM CaCl_2_, 5 mM KCl, 2 mM MES, pH 5.7) was added. The protoplast solution was centrifuged at 250Xg for 5 minutes. Number of protoplast was adjusted to 10^6^/ml using W5 solution and incubated on ice for 30 minutes. Protoplasts were again centrifuged at 250X g for 2 minutes and suspended in MMg solution (0.6 M mannitol, 15 mM MgCl_2_, 4 mM MES, pH 5.7) to obtain 10^6^ cells (Lee et al. 2008).

#### Transient transfection of *Nicotiana tabacum* By-2 cell line

Constructs of effector plasmid, 9 μg (Camv35S: bZIP10, Camv35S: bZIP25, Camv35S: bZIP53, and Camv35S:A-ZIP53), reporter plasmid, 9 μg (2S2:GUS), and normalization vector, 1μg (CaMV35S: NAN) were mixed with 100 μl of protoplast (2X10^4^ protoplasts) and mixed gently. 110 μl of PEG solution (Prepare 20–40% (wt/vol) PEG3500 in ddH_2_O containing 0.2 M mannitol and 100 mM CaCl_2_) was added. Transfection mixture was incubated at room temperature for 15 minutes. Mixture was diluted with 400-440 μl of W5 solution, gently inverted to stop the transfection process. Protoplasts were resuspended with 1 ml of WI solution (4 mM MES – pH 5.7, 0.5 M mannitol, and 20 mM NaCl, it can be stored at room temperature) and divided into six different tube. Protoplasts were incubated in dark at room temperature (20 ᴼC – 25 ᴼC) for 16 –18 hours. Protoplast suspension was centrifuged at 100x g for 2 minutes at room temperature. Supernatant was removed and samples were kept in -80 ᴼC for further analysis (Yoo et al. 2007). GUS/NAN ratio was quantified as described earlier (Alonso et al. 2009).

#### Protoplast isolation and transient transfection using *Arabidopsis* leaves

Protoplasts were isolated as described earlier (Jain et al. 2017). Four weeks old Col-0 plants with well expanded healthy leaves were selected for protoplast isolation. 0.5 mm – 1 mm part of leaves were cut from the middle with sharp razor blade. For 10^6^ - 10^7^ / gm fresh protoplast approximately 40 - 50 leaves are required. Cut sections were digested in 5 – 10 ml of enzyme solution {20 mM MES (pH 5.7) containing 1.5% (wt/vol) cellulase (sigma), 0.4% (wt/vol) macerozyme (sigma), 0.4 M mannitol, and 20 mM KCl. Solution was heated at 55 ᴼC for 10 minutes to inactivate the DNAse and protease. Cool it at room temperature and add 10 mM CaCl_2_, 1–5 mM β-mercaptoethanol (optional), and 0.1% BSA (Final enzyme solution should be clear light brown). Leaves were vaccum infilterated with the final enzyme solution for 30 minutes in dark using desiccator. Digestion was continued by putting the leaves in dark for 3 -4 hours at room temperature. The green color of solution resembles the release of protoplasts. Number of released protoplasts were checked under the microscope. Enzyme solution was diluted with equal volume of W5 solution (2 mM MES (pH 5.7) containing 154 mM NaCl, 125 mM CaCl_2_, and 5 mM KCl) that can be stored at room temperature. Enzyme solution was filtered with muslin cloth and centrifuged at 100 g in a 30 ml round bottomed tube for 1 minute. Supernatant was removed and pellet was re suspended at 2 × 10^5^ cells in W5 solution. Protoplast were kept in ice for 30 minutes and allowed to settle. W5 solution was removed without disturbing the settled protoplast. Protoplast were resuspended at 2 × 10^5^ /ml in MMG solution {4 mM MES (pH 5.7) containing 0.4 M mannitol and 15 mM MgCl_2_. The prepared MMG solution can be stored at room temperature (Yoo et al. 2007).

To confirm the heterodimeric interaction between the *A-ZIP53* and target proteins, constructs of effector, reporter, and normalization vector were transformed and subjected for the GUS-NAN activity as described earlier (Alonso et al. 2009; Jain et al. 2017).

### RNA extraction and Illumina sequencing

Total RNA was extracted from the wild type and *A-ZIP53* expressing *Arabidopsis* using ZR plant RNA miniprep (ZYMO Research) as per manufactures instruction. The quality and quantity of RNA was checked on 1 % denaturing RNA agarose gel and NanoDrop/Qubit fluorometer, respectively. The RNA-seq paired end sequencing library were prepared from the QC passed RNA samples using illumina TrueSeq stranded mRNA sample preparation kit. Briefly, mRNA was enriched from the total RNA using poly-T attached magnetic beads, followed by enzymatic fragmentation, 1^st^ strand cDNA conversion using Superscript II and Act-D mix to facilitate RNA dependent synthesis. The 1^st^ strand cDNA was then synthesized to second strand using second strand mix. The ds cDNA was then purified using Ampure XP beads followed by A-tailing, adapter ligation, and then enriched by limited number of PCR cycles.

### Cluster generation and Sequencing

After obtaining the Qubit concentration for the libraries and the mean peak size from Agilent Tape Station profile, the PE illumine libraries were loaded onto NextSeq 500 for cluster generation and sequencing. Paired-End sequencing allows the template fragments to be sequenced in both the forward and reverse directions on NextSeq500. The adaptors were designed to allow selective cleavage of forward strand after re-synthesis of reverse strand during sequencing. The copied reverse strand will then use to sequence from the opposite end of the fragment.

### RNA seq analysis

Adaptor trimming and quality trimming of the samples (wild-type and three biological replicates of the A-ZIP53 expressing transgenic) were performed using Trimmomatic- 0.35. The sequenced raw data was processed to obtain high quality clean reads using Trimmomatic to remove adaptor sequences, ambiguous reads (reads with unknow nucleotides “N” larger than 5%) and low-quality sequences (read with more than 10 % quality threshold (QV) <20 phred score). A minimum length of 50 nucleotide (after trimming) was applied. After removing the adaptor and low-quality sequences from the raw data, high quality sequences were obtained. This high quality paired-end reads were used for the referenced based read mapping. The high-quality reads were mapped on the reference genome of *Arabidopsis thaliana* using TopHat v2.1.1 with the default parameters.

### Gene ontology (Go) and differential gene expression (DGE) analysis

The DGE was carried out using cutdiff v1.3.0. Fold change (FC) values greater than zero wer considered as upregulated whereas less than zero as downregulated. P value threshold of 0.05 was used to filter statistically significant results. For GO analysis Singular Enrichment Analysis (SEA) of agri GO(http://bioinfo.cau.edu.cn/agriGO/analysis.php) was used. Hypergeometric tests with Hochberg FDRs (false discovery rates) were performed using the default parameters to adjust the P-value <0.05 for obtaining significant GO terms.

### Quantitative real-time PCR (qRT-PCR)

The differential gene expression was validated by qRT-PCR. Total RNA (2 μg) was isolated (Spectrum Plant Total RNA kit) from the leaves (T1 generation) and immature siliques (T2 and T3 generation) from wild-type and A-ZIP53 expressing *Arabidopsis*. Contamination of genomic DNA was removed by the Turbo DNA-free kit (Invitrogen, ThermoFisher, USA).

Later, cDNA was synthesized (Invitrogen Superscript® III Reverse Transcriptase) as per manufacturer’s protocol. For qRT-PCR gene specific primers were used. The template concentration was 10-15 ng while the concentration of forward and reverse primer was 10 ng in SYBR select Master Mix (ABI). The PCR was performed using the ABI 7700 sequence detector (Applied Biosystems, USA) as per manufacturers instruction. Two biological and three technical experiments were taken for each experiment. Statistical analysis was done using Origin 6.1.

### Total protein extraction and Western blotting

Total protein extract was prepared by grinding the 5 days old seedling grown in the long day conditions using the protein extraction buffer (50 mm Tris–HCl, pH 8, 150 mm NaCl, 1 mm EDTA, pH 8.0, 1% SDS, 1 mM PMSF, 1 mM DTT, protease inhibitor) (Mair, 2015). Total protein was quantified using the bradford assay method (Bradford 1976). Protein was separated on the 10 % SDS PAGE and transferred on the polyvinylidene difluoride (PVDF) membranes (Hybond™-P; GE Healthcare, Piscataway, NJ, USA) using the wet transfer method (1 hour, 100 volt, 4 °C). Transferred blot was incubated in the blocking buffer (3 % skimmed milk in TBST buffer (20 mm Tris-HCl, pH 7.5, 200 mm NaCl and 0.1 % Tween 20) for 1 hour at the room temperature to inhibit the non-specific antibody interactions. For immunoblotting of *A-ZIP53* the blocked blot was incubated with the 1:5000 dilution of HRP labeled rat anti-T7 antibody (Thermo Scientific) at 4 °C overnight. The fluorescence signals were detected by the fluorescence imager using the SuperSignal^TM^ West Pico PLUS Chemiluminescent Substrate (Thermo Scientific).

### Co-Immunoprecipitation of proteins in Transgenic *Arabidopsis*

Pull-down assay was used to extract the target proteins from mixture (cell lysate). For Co-IP of *A-ZIP53* from the protein extract of transgenic plants, antibodies targeting T7 tag were mixed with protein lysate in the protein extraction buffer. Mixture was kept at 4 °C for 2 hours. “Protein extracted” with T7 tag antibodies were incubated with protein A containing magnetic beads (Dyna beads, Thermo fisher) for 2 hours at 4 °C. Beads with immobilized antibodies were collected through magnetic separation rack (NEB). Beads were washed with protein extraction buffer for 1 minute, washed beads were collected with magnets and washing was repeated. Final washing was done with autoclave water for 1 minute. Beads with immobilized protein of interest were again collected and resuspended in 1× Laemmli sample buffer (32.9 mM Tris HCl (pH 6.8), 13.15 % glycerol, 1 % SDS, 0.01 % bromophenol blue, and 355 mM β mercaptoethanol), boiled at 95 °C for 5 minutes and centrifuged. Beads were collected with magnet and supernatant was separated on 15 % SDS PAGE.

### In-Gel digestion of immunoprecipitated proteins

To check the identity of protein with mass spectrometry, the Coomassie stained gel was rinsed with water and bands were excised with the clean scalpel. Excised bands were chopped with the scalpel and washed with 40 mM ammonium bicarbonate (ABC, pH 8.5), and dried with the acetonitrile (ACN). Bands were gently agitated and again washed with the destaining solution for complete destaining. Destained gel pieces were incubated in 0.5 ml of 100 % ACN for 10-15 minutes until gel pieces shrunk and become opaque. The liquid was discarded and gel pieces were incubated in the reduction solution (5 mM DTT, 40 mM ABC) at 60 °C for 5 minutes. DTT was washed off and the cystein were alkylated using the alkylation solution (20 mM IAA in 40 mM ABC) for 10 minutes in dark at room temperature. Gel pieces were dehydrated with 100 % ACN and trypsinized using the sequencing grade trypsin (Sigma Chemical Co, USA) at a concentration of 10 ng/μl in 40 mM ABC and incubated overnight at 37 °C. The digestion was stopped by the addition of approximately 0.1 % of the formic acid with a incubation at 37 °C for 10 minutes. Tubes were spun and the supernatant was incubated with 100 μl of extraction buffer (5 % formic acid, 40 % ACN) for 10 minutes at room temperature. Tubes were again centrifuged and the supernatant was mixed with extraction buffer. The final extraction was carried out with the 100 % ACN by incubating at room temperature for 10 minutes. The supernatant was collected and dried using Speed Vac (Thermo fisher Scientific, USA). Trypsinized peptides were sequenced using the Triple TOF M 5600 mass spectrometer (AB Sciex Pte. Ltd., USA) attached with the Nano-LC (Eksigent Technologies Llc, USA). The dried trypsinized 0.1 μg of protein were resuspended in 0.1 % formic acid. 10 μl of the sample was passed through the desalting trap column (Chrom-XP, C-18-CL-3μM, 350 μM X 0.5mm, Eksigent Technologies LLC., USA). After desalting peptides were separated on C18 matrix (3C-18-CL-120, 3 μM, 120A, 0.075 × 150 mm, Eksigent Technologies LLC., USA) before peptide sequencing. The eluents used were eluent A, degassed ionized water with 0.1 % (v/v) formic acid, eluent B, 100 % acetonitrile (containing 0.1 % (v/v) formic acid with a linear gradient of 5-95 % for 120 mins with a flow rate of 300 nl/min. After separation peptides were subject to tandem mass (MS/MS) analysis. The Nano-LC was directly coupled to a Triple TOFM 5600 mass spectrometer, which was operated in an information dependent acquisition (IDA) mode. For IDA one full scan (m/z 350-1250) was followed by 8 MS/MS scans and the electrospray voltage was set to 2500 V.

### 3.2.64 Data interpretation

Raw spectra for the peptide identification were interpreted using Protein Pilot 4.0 (ABsciex). The peptide spectra were searched against the *Arabidopsis thaliana* entries using uniprot database under following parameters: the peptide tolerance was set to 1 Da and MS/MS was set to 0.8 Da. Trypsin was selected as a protease.

## ACKNOWLEDGMENTS

Authors are grateful to Prof. Wolfgang Dr**Ö**ge-Laser, University of Würzburg, Germany for providing overexpressing full-length *bZIP53*, *bZIP10*, *bZIP25*, *bZIP14*, *bZIP17*, *bZIP19*, *bZIP29*, *bZIP34*, *bZIP69*, *NAN*, and *GUS* reporter gene plasmids. We thank Executive Director, NABI, Department of Biotechnology (DBT), and GOI for providing research facilities. PJ acknowledges the financial assistance in the form of JRF and SRF from DBT, GOI. We thank Shrikant Mantri and Gazaldeep Kaur for their help in proteome and transcriptome data analysis. This work is supported by the core grant of National Agri-Food Biotechnology Institute, Government of India (GOI).

## AUTHOR CONTRIBUTION

**Conceptualization**: Prateek Jain and Vikas Rishi.

**Data curation**: Prateek Jain and Vikas Rishi.

**Formal analysis**: Prateek Jain.

**Methodology**: Prateek Jain.

**Project administration**: Prateek Jain and Vikas Rishi.

**Software**: Prateek Jain.

**Validation**: Prateek Jain.

**Writing:** original draft: Prateek Jain.

**Writing – review & editing:** Prateek Jain and Vikas Rishi.

## Supplementary Figure

**Supplementary FigureS1.**
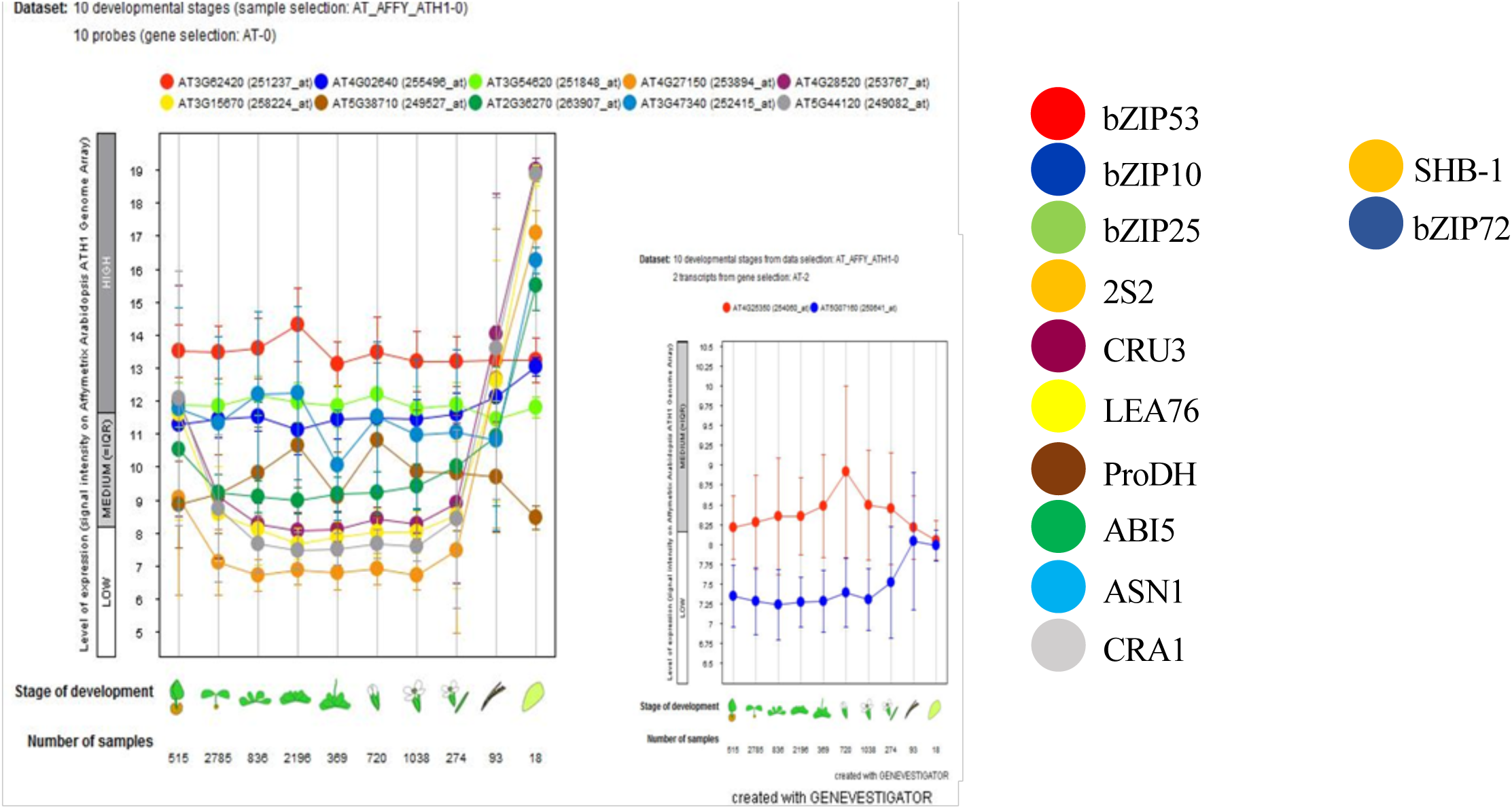
Geneinvestigator revealed the expression of bZIP53 (Red), bZIP10 (Blue–first panel), bZIP25 (Light green), target genes (**2S2:** Orange, **CRU3:** Violet, **LEA76:** Yellow, **ProDH:** Brown, **AS**^1 m^**N**^m^**1:** Sky Blue, **CRA1**^1^:^mm^G^1^r^m^e^m^y), Non heterodimerizing partner (^1^**b**^mm^**ZIP72**: Blue-second panel, **bZIP39**: Green) and Non target gene (**SHB1**: Orange-second panel) in the different development stage of *Arabidopsis*.

**Supplementary FigureS2.**
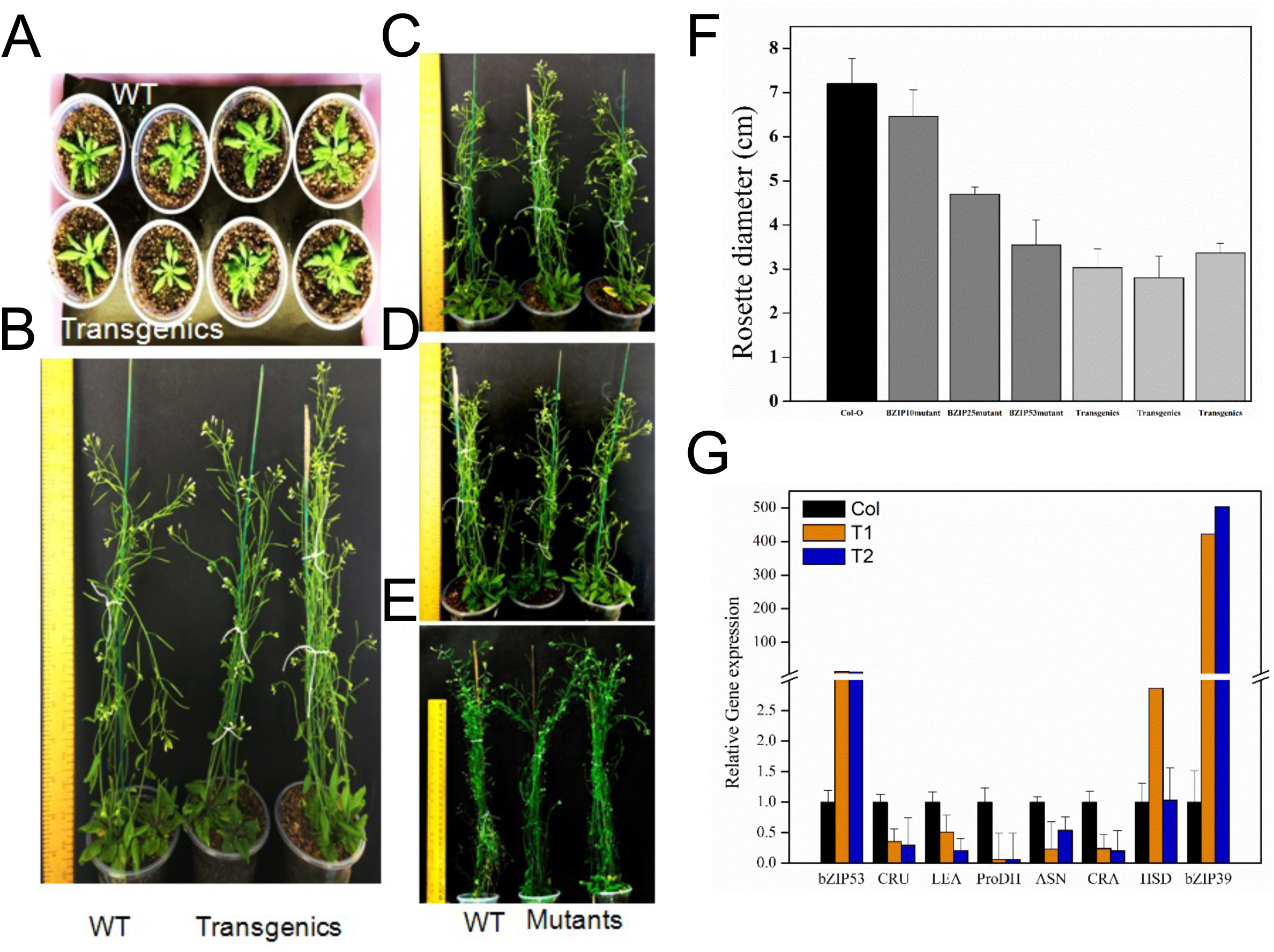
Phenotypic alteration and expression analysis of target genes of bZIP53 and its dimerizing partners in T3 generation. **(A**1 m,m**B)** Growth of the four an1 mdm eight week old wild type and transgenic **(C, D, E)** Eight weeks old mutants of *bzip10*, *bzip25*, and *bZIP53* **(F)** Differences in the rosette diameter of three week old transgenic, mutants (*bZIP53*, *bzip1*0, and *bzip25*) and wild-type under standard condition. Error bar represent ± mean and SD of 8-10 individual plants**. (G)** Expression of target genes of *bZIP53*, *bZIP10*, and *bZIP25* involved in the seed maturation from the immature silique and seeds of transgenic. Error bar represent ±S.D. of three technical replicates.

**Supplementary FigureS3.**
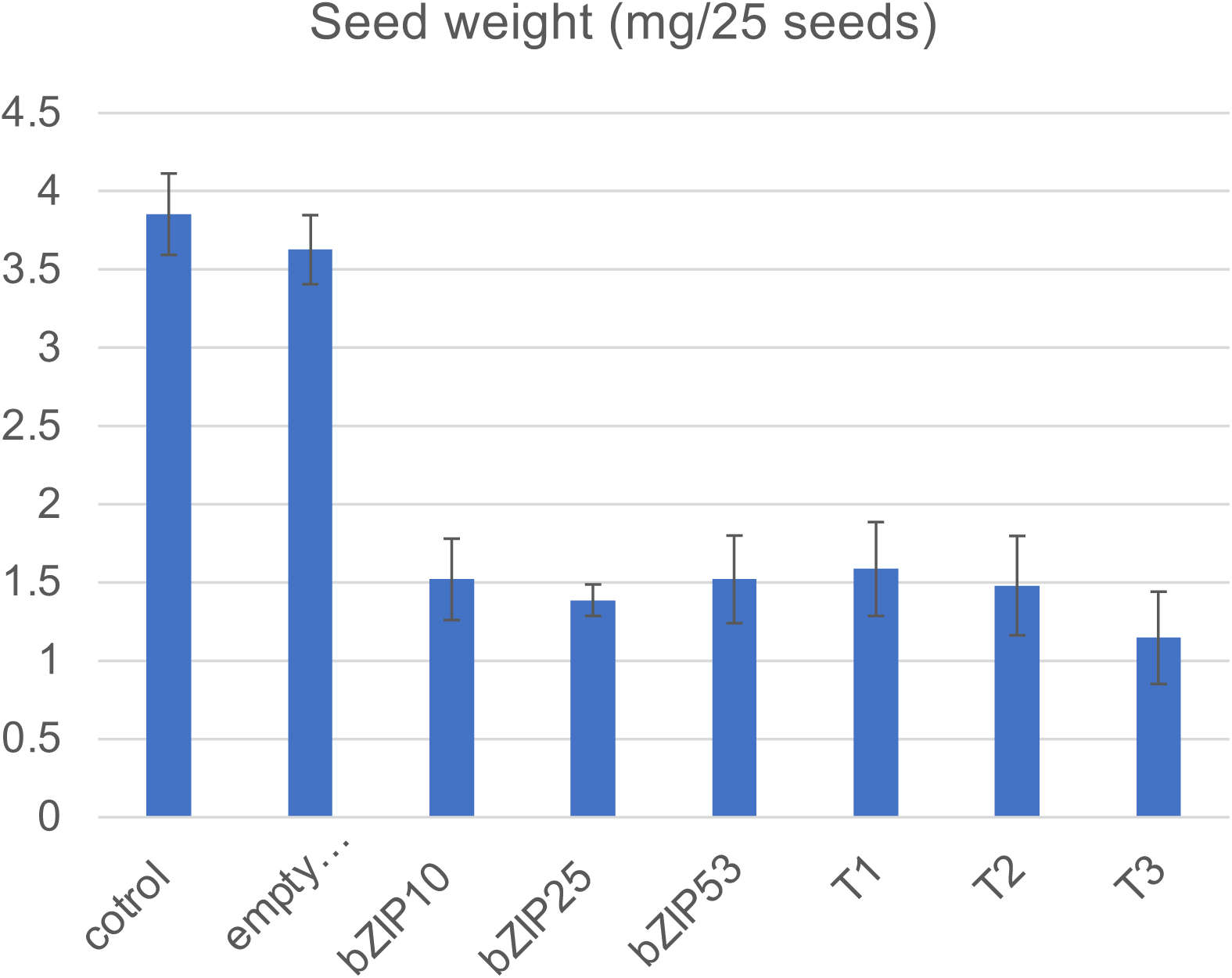
Seed weight calculated (mg/25 seeds). Transgenic has lesser weight compare to WT. ±Mean and S.D. of three independent replicate was calculated. One way Anova was used to check the level of significance (p<.01, ns = not significant).

**Supplementary Figure S4.**
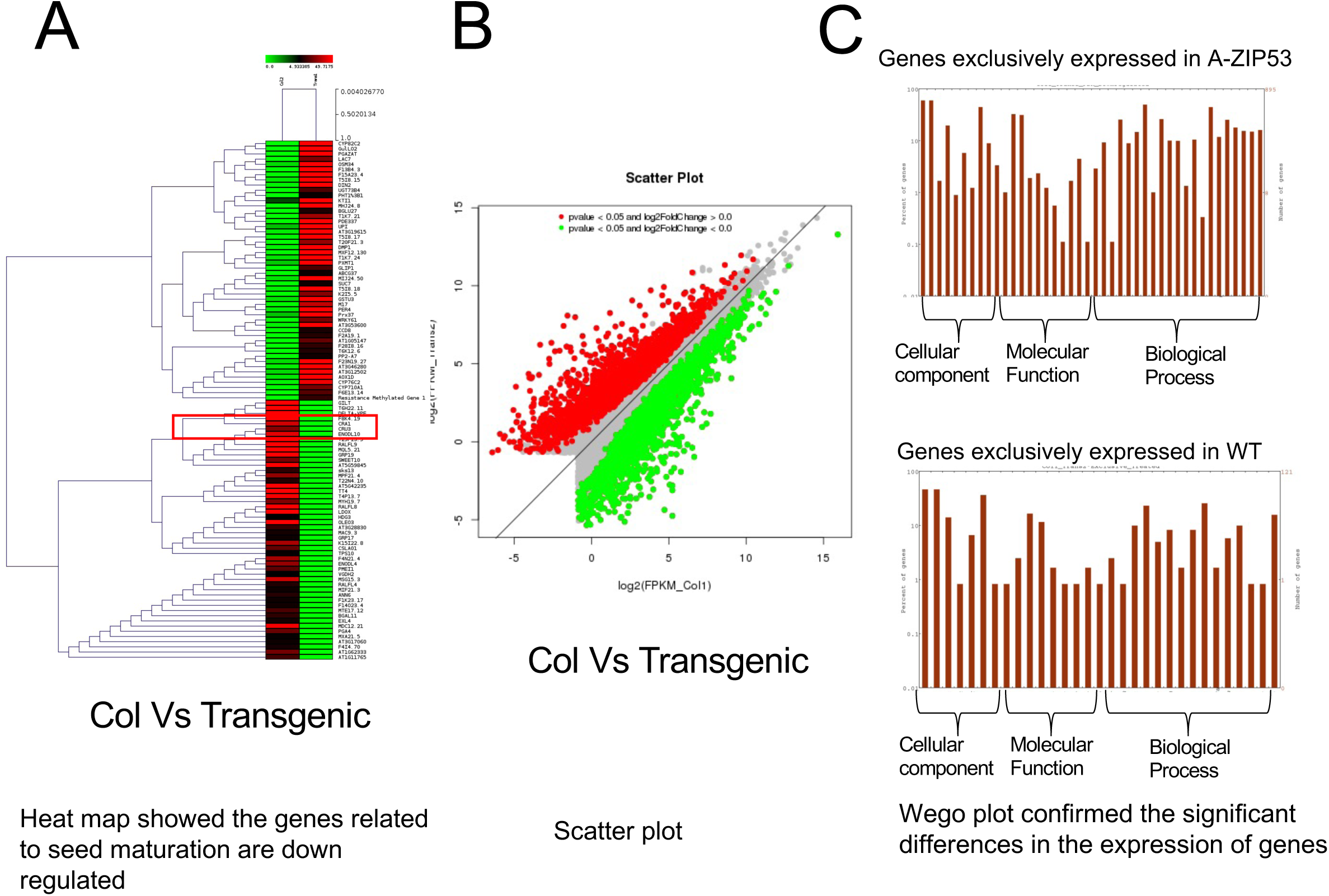
Peptide fingerprinting of immunoprecipitated leucine zipper domain of bZIP53/A-ZIP53.

**Supplementary FigureS5.**
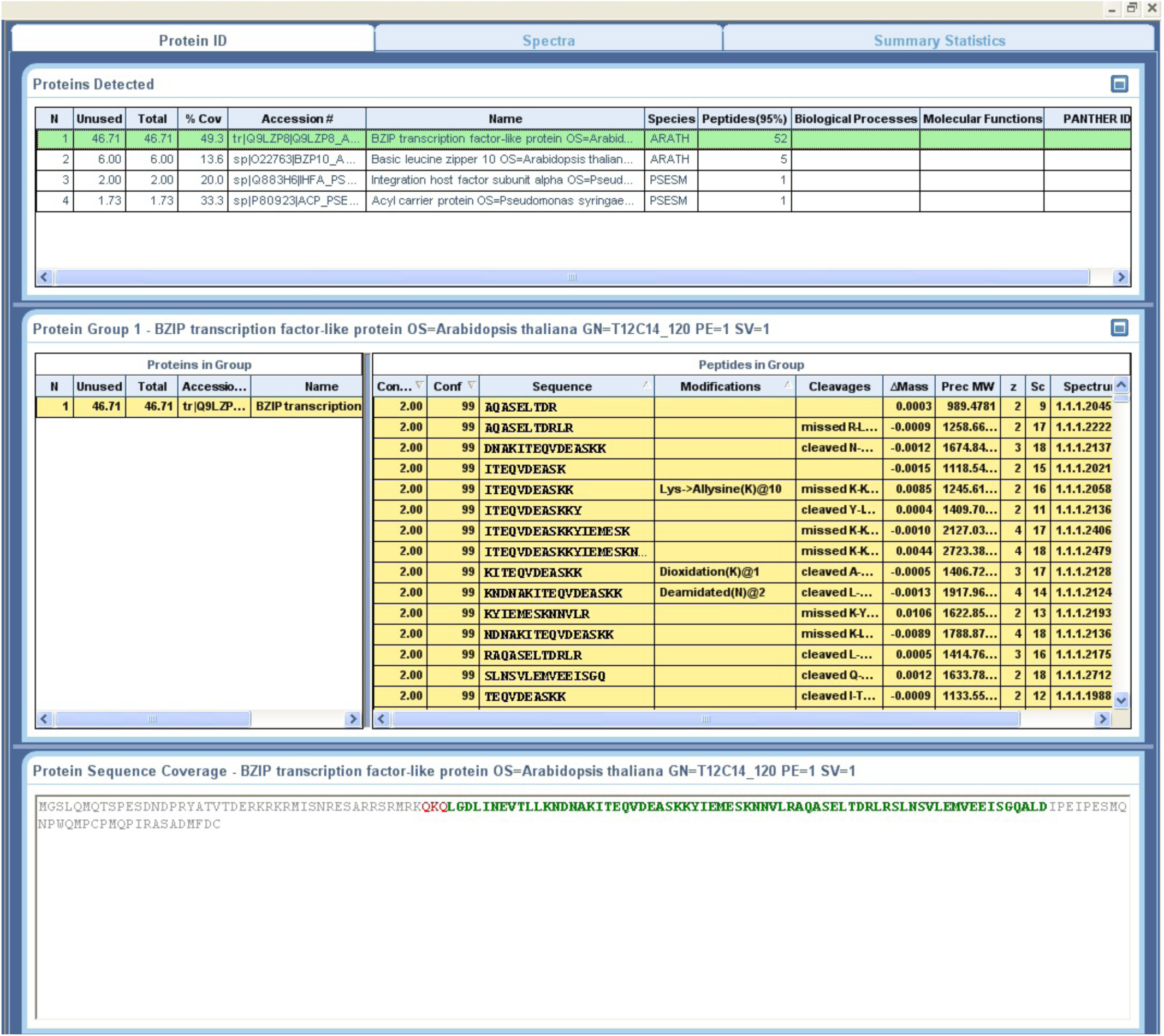
Peptide fingerprinting of Immunoprecipitated leucine zipper domain of bZIP53/A-ZIP53 from total protein soup of Arabidopsis.

**Supplementary Figure S6.**
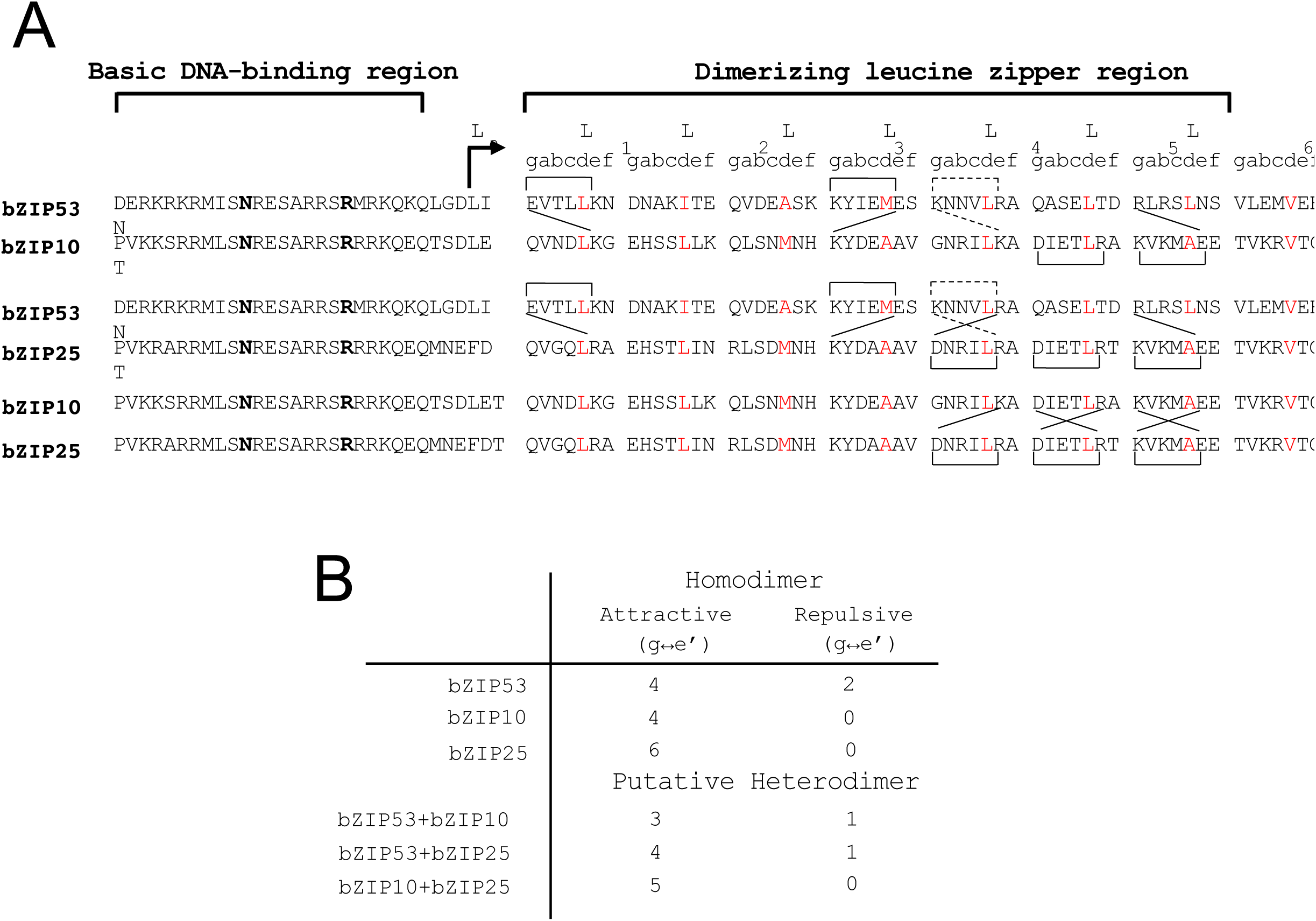
Amino acid sequences of bZIP53, bZIP10, and bZIP25. **A)** The delineation of N-terminal basic DNA-binding region followed by dimerizing leucine zipper region. Amino acid sequences represented by the single-letter code are aligned with respect to an invariant asparagine (N) and arginine (R) (shown in bold) in the basic region. Only ten amino acids upstream of asparagine are shown. Tenth amino acid (typically a leucine; L_o_) from invariable arginine in the basic DNA-binding region marks the start of the dimerizing leucine zipper. The leucine zipper sequence is grouped into heptad **(a,b,c,d,e,f,g)_n=8_**. The limit of a coiled coil at C-terminus is defined by the presence of a proline or two consecutive glycines, both likely helix-breaking residues and the absence of charged amino acids in g and e′ positions in a heptad In homodimer coiled coil, interhelical interactions between amino acids in the g position with those in the following e′ position are shown as square brackets. Solid square brackets depict attractive interactions between amino acids with opposite charges in g and e′ positions (E↔K, K↔E, D↔R) whereas interhelical repulsive interactions between g and e′ position amino acids are shown by discontinuous square brackets (K↔R). In the putative heterodimer coiled coil (bZIP53+bZIP10, bZIP53+bZIP25 and bZIP25+bZIP10) attractive interactions between amino acids at g and e′ position are shown by solid diagonal lines (E↔K, R↔E, R↔D, K↔D) and repulsive interactions are depicted by discontinuous lines (K↔K,K↔R). B) The number of attractive and repulsive g↔e′ salt bridges formed in three homodimers and three putative heterodimers.

**Supplementary Figure S7.**
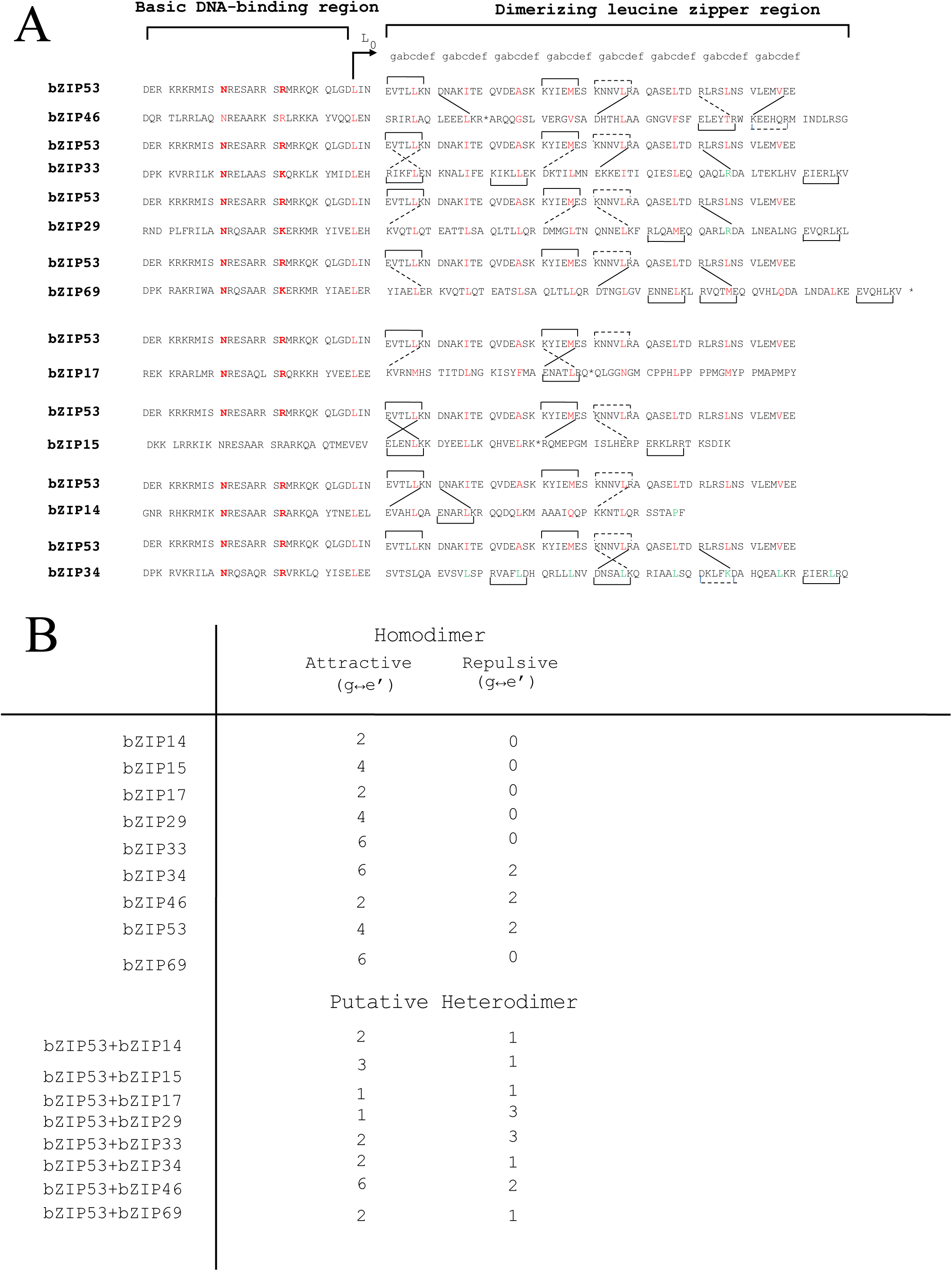
**Supplementary Figure S6** Amino acid sequences of B-ZIP53, B-ZIP14, B-ZIP15, B-ZIP17, B-ZIP29, B-ZIP33, B-ZIP34, B-ZIP46, and B-ZIP69. **A**) At the top is the delineation of N-terminal basic DNA-binding region followed by dimerizing leucine zipper region. Amino acid sequences represented by the single-letter code are aligned with respect to an invariant asparagine (N) and arginine (R) (shown in red) in the basic region. Only ten amino acids upstream of asparagine are shown. Tenth amino acid (typically a leucine; Lo) from invariable arginine in the basic DNA-binding region marks the start of the dimerizing leucine zipper. The leucine zipper sequence is grouped into heptad (**a,b,c,d,e,f, g**)_n=8_. The limit of a coiled coil at C-terminus is defined by the presence of a proline or two consecutive glycines, both likely helix-breaking residues and the absence of charged amino acids in **g** and **e’** positions in a heptad. Also shown below is a consensus sequence for B-ZIP motif, where Ψ represents any hydrophobic amino acid. Proteins are placed in three groups with three homodimers and three potential heterodimers. In homodimer coiled coil, interhelical interactions between amino acids in the **g** position with those in the following **e’** position are shown as square brackets. Solid square brackets depict attractive interactions between amino acids with opposite charges in **g** and **e’** positions (E↔K, K↔E, D↔R) whereas interhelical repulsive interactions between **g** and **e’** position amino acids are shown by discontinuous square brackets (K↔R). In the putative heterodimer coiled coil (B-ZIP53+B-ZIP14, B-ZIP53+B-ZIP15, B-ZIP53+B-ZIP29, B-ZIP53+B-ZIP33, B-ZIP53+B-ZIP34, B-ZIP53+B-ZIP46, B-ZIP53+B-ZIP69) attractive interactions between amino acids at **g** and **e’** position are shown by solid diagonal lines (E↔K, R↔E, R↔D, K↔D) and repulsive interactions are depicted by discontinuous lines (K↔K,K↔R). B) The number of attractive and repulsive **g↔e’** salt bridges formed in three homodimers and three putative heterodimers. **B**) The number of attractive and repulsive **g↔e’** salt bridges formed in homodimers and putative heterodimers

**Supplementary Figure S8.**
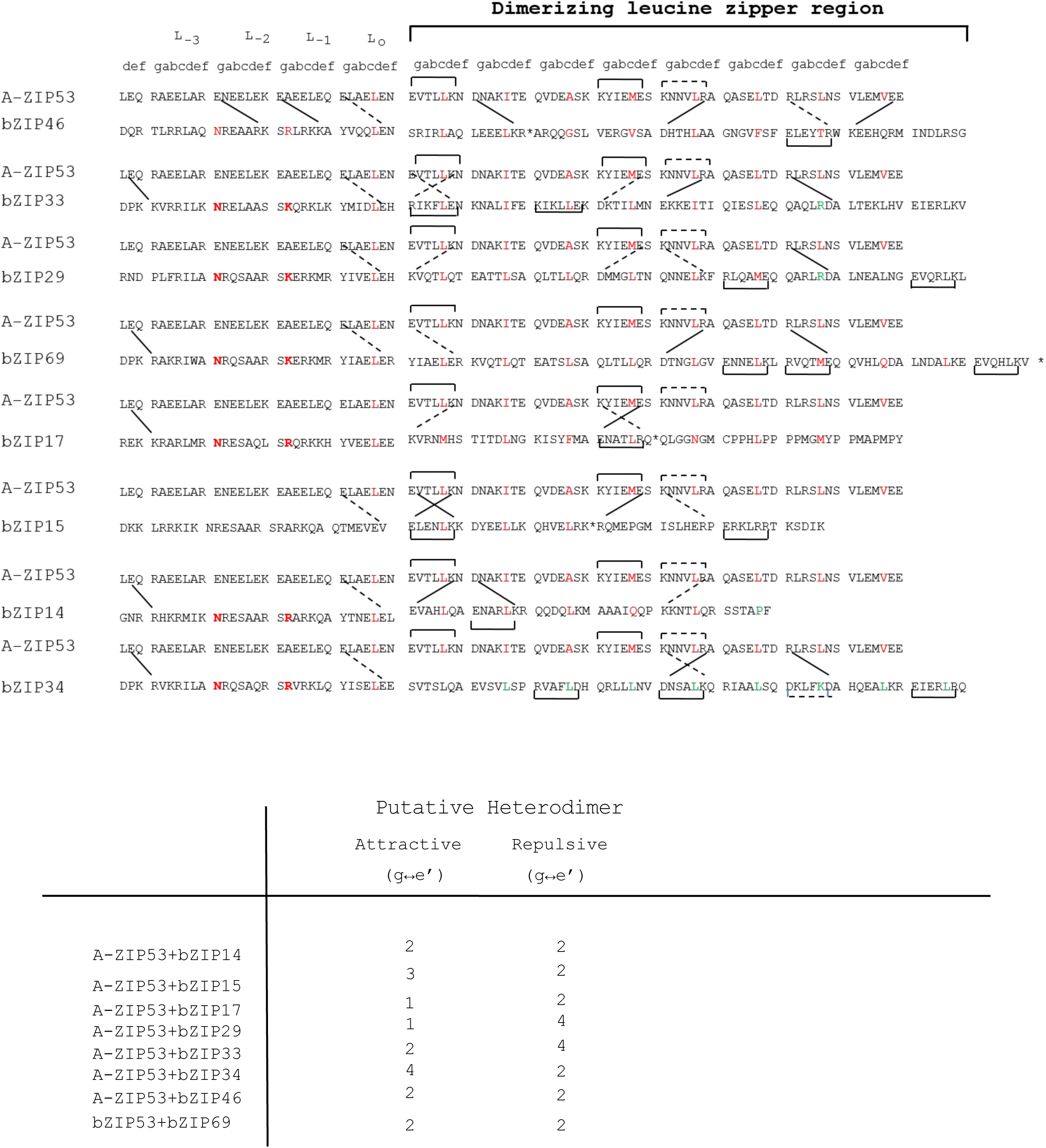
Heterodimers between A-ZIP53 and the immunoprecipitated B-ZIPs **A)** Alignment of acidic helical extensions used in this study with DNA-binding region of wild type B-ZIPs showing the possible interhelical interactions in homodimers and heterodimers. At the top is shown the coiled coil heptad designations (**a,b,c,d,e,f,** and **g**). A-ZIP53 acidic extension are aligned with the basic region of B-ZIPs. invariable asparagine at **g** position (L_-2_), and arginine at **a** position (L_-3_) are shown in bold. In homodimer coiledcoil, interhelical interactions between amino acids in the **g** position with those in the following **e’** position are shown as square brackets. Solid square brackets depict attractive interactions between oppositely charged amino acids in **g** and **e’** positions whereas broken square brackets shows repulsive interactions due to the presence of similar charges (E↔E). In the putative heterodimeric coiled-coil, attractive interactions between **g** and **e’** positions amino acids are shown by solid diagonal lines. **B**) The number of attractive and repulsive **g↔e’** salt bridges formed in homodimers and putative heterodimers.

## Supplementary Tables

**Table 1** DNA binding sites of target bZIPs on their corresponding genes.

**Table 2** Primer sequences for the qRT PCR.

**Table 3** Primer sequence for the cloning of A-ZIP53 into the pRI101AN

**Table 4** Comparisons of length and width of seeds

**Table 5** Dry weight of mature seed (25 seeds)

**Table 6** Differences in the flower size of wildtype, mutant (*bZIP53, bzip25, and bzip10*) and transgenic

**Table 7 Differences in the length and width of siliques of** wildtype, mutants (*bZIP53, bzip25, and bzip10*), and transgenic.

Table 8 Genes downregulated in A-ZIP53 expressing transgenic plants

Table 9 Immunoprecipitated peptides in A-ZIP53 epressing transgenic plants

