## Supplementary material for "Synergistic regulation of bZIP53 and dimerizing partners results in abnormal seed phenotype in Arabidopsis: Use of a designed dominant negative protein A-ZIP53": Table

**Supplementary Table 1 Probable DNA binding site of target bZIP transcription factor**

| **S.No.** | **bZIP** | **Gene** | **Binding site** |
|---|---|---|---|
| 1. | bZIP10 | CRU | 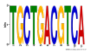 |
| 2. | bZIP25 | CRU | 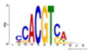 |
| 3. | bZIP39 | CRU | 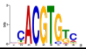 |
| 4. | bZIP10, bZIP25 | CRU | 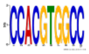 |
| 5. | bZIP10, bZIP25 | CRU | 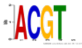 |
| 6. | bZIP10, bZIP25 | LEA | 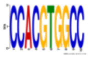 |
| 7. | bZIP10, bZIP25 | LEA | 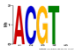 |
| 8. | bZIP25 | LEA | 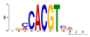 |
| 9. | bZIP39 | LEA | 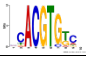 |
| 10. | bZIP10, bZIP25 | ASN | 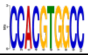 |
| 11. | bZIP10, bZIP25 | ASN | 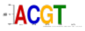 |
| 12. | bZIP25 | ASN | 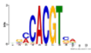 |
| 13. | bZIP39 | ASN | 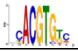 |
| 14. | bZIP10, bZIP25 | HSD | 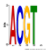 |
| 15. | bZIP25 | HSD | 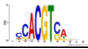 |
| 16. | bZIP39 | HSD | 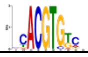 |
| 17. | bZIP10, bZIP25 | ProDH | 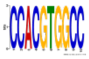 |
| 18. | bZIP25 | ProDH | 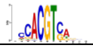 |
| 19. | bZIP39 | ProDH | 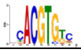 |
| 20. | bZIP25 | CRA | 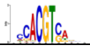 |
| 21. | bZIP25 | CRA | 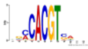 |
| 22. | bZIP25 | CRA | 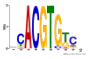 |
| 23 | bZIP | | |

**Supplementary Table S2** Primer for qRT PCR

| **Gene** | **Primer** | **Sequence** |
|---|---|---|

| | | |
|---|---|---|
| A-ZIP53 | Forward<br>Reverse | CTG GAA CAA CGT GCT GAG GAA CTG<br>GTT ATC ATT CTT GAG AAG AGA GAC |
| UBI | Forward<br>Reverse | GCTCTTATCAAAGGACCTTCGG<br>CGAACTTGAGGAGGTTGCAAAG |
| bZIP53 | Forward<br>Reverse | TAATGATCCGAGGTACGCCAC<br>TGCTTCTGTTTCCTCATCCTTG |
| CRU | Forward<br>Reverse | TAGATGTTCTCCAAGCCACCG<br>AACGGAAACACCAACACATCG |
| 2S2 | Forward<br>Reverse | ATTTGCAAGATCCAGCAAGTTG<br>AATACATTTAGCCTCAAACATC |
| LEA76 | Forward<br>Reverse | ACAAAGAGCATTATCCAGGAAGT<br>ACACAAAGATACTTTCATATCGT |
| ProDH | Forward<br>Reverse | GCATCAAACGGTTCTGGTTTC<br>TGTTTATCGATCCCGAGGTCA |
| ASN1 | Forward<br>Reverse | TTCAACGCCTTATGAGCCTCTT<br>CACCAGAGAGCAAAACTCCAAA |
| CRA1 | Forward<br>Reverse | AGCCCAAATCCAGATCGTAAAC<br>TCACCACCGAGAAACCTTGTG |
| HSD1 | Forward<br>Reverse | TGCCGGAAACAAAGATACGTG<br>AGTAACCGACAACCCCACTCA |
| bZIP39 | Forward<br>Reverse | GAT CGA TAT GGG AAA CAG CAT GCT<br>GAT CCC ACA AGC TTG CTA TCC A |
| SHB1 | Forward<br>Reverse | AAGAATGGTGGAAGACAGAGAT<br>GAGAAGCAGCAACGATGGT |

**Supplementary Table S3** Primer sequence to amplify A-ZIP53 for the cloning into pRI-101AN

| **Primer** | **Sequence** |
|---|---|
| Forward | GGGAGACCACAACGGTTTCCCTC |
| Reverse | **GACGGCGAATTCTAATCCAAAGCCTGACCACTAA** |

**Supplementary Table S4 Comparisons of length and width of seeds**

| S.No. | Genotype | Length | S.D. | Width | S.D. |
|---|---|---|---|---|---|
| **1.** | Col-O | 539.4735 | 91.2474 | 313.0386 | 60.57069 |
| **2.** | Empty plasmid | 523.3212 | 92.47669 | 310.4526 | 87.76217 |
| **3.** | *bzip53* | 365.9634 | 54.10735 | 187.1833 | 58.74303 |
| **4.** | *bzip10* | 439.5349 | 91.21795 | 175.1596 | 84.42437 |
| **5.** | *bzip25* | 416.207 | 77.34801 | 131.2121 | 72.57546 |
| **6.** | L-1 | 286.7765 | 70.95585 | 169.0514 | 43.5315 |
| **7.** | L-2 | 359.0172 | 54.60866 | 178.1991 | 70.91795 |

Mature seed were analysed. Mean value and standard deviation of triplicate analyses are presented. Level of significance was checked using T-test (one-tailed, two sample with equal variance) Transgenic was ** significantly different from WT ($p<.01$).

**Supplementary Table S5 Dry weight of mature seed (25 seeds)**

| | Control | *bZIP53* | bZIP10 | bZIP25 | Transgenic | Transgenic |
|---|---|---|---|---|---|---|
| | 0.85 | 0.288 | 0.31 | 0.32 | 0.17 | 0.48 |
| | 0.91 | 0.24 | 0.45 | 0.37 | 0.26 | 0.68 |
| | 0.96 | 0.3 | 0.38 | 0.35 | 0.19 | 0.36 |
| Average | 0.906667 | 0.276 | 0.38 | 0.346666667 | 0.506667 | 0.206666667 |
| Stdev. | 0.055076 | 0.031749016 | 0.07 | 0.025166115 | 0.161658 | 0.047258156 |
| T-test | | 4.00369E-05 | 0.000226 | 5.30749E-05 | 4.14762E-05 | 0.005252 |

Dry and mature seeds were used. Mean and standard deviation from three replicates (N =25). Level of significance was compared between WT and transgenic. Level of significance was checked using T-test (one-tailed, two sample with equal variance) one way Anova. Transgenic was ** significantly different from WT ($p<.01$).

**Supplementary Table S6 Differences in the flower size of wild type, mutant, and transgenic**

| S.No. | Genotype | Height | ± S.D. (n =10-15) | Width | --± S.D. (n =10-15) |
|---|---|---|---|---|---|
| **1.** | Col -O | 3.44 | 0.24194 | 1.7846 | 0.13625 |
| **2.** | *bzip10* | 2.951 | 0.27225 | 1.33475 | 0.17642 |
| **3.** | *bzip25* | 2.689 | 0.15235 | 1.3694 | 0.15494 |
| **4.** | *bZIP53* | 1.954 | 0.18758 | 1.078 | 0.12921 |
| **5.** | L1 | 1.885 | 0.22887 | 0.9888 | 0.12272 |
| **6.** | L2 | 1.828 | 0.1362 | 1.0923 | 0.14968 |
| **7.** | L3 | 1.768 | 0.18796 | 1.0948 | 0.19545 |

Flowers were taken from six week old plants. Mean value and standard deviation from three replicates (N = 10-15). Level of significance was compared between WT and transgenic using T-test (one-tailed, two sample with equal variance) was used for the statistical analysis for three replicates. **Transgenics were ** significantly different from WT (p<.01).**

**Supplementary Table S7** Differences in the length and width of siliques of wild-type, mutants, and transgenic

| | **Col O** | | **L1** | | **L2** | | ***bZIP25*** | | ***bZIP10*** | | ***bZIP53*** | |
|---|---|---|---|---|---|---|---|---|---|---|---|---|
| | Length (mm) | Width (mm) | Length (mm) | Width (mm) | Length (mm) | Width (mm) | Length (mm) | Width (mm) | Length (mm) | Width (mm) | Length (mm) | Width (mm) |
| **1** | 14.46 | 0.86 | 8.9 | 0.9 | 6.14 | 0.78 | 13.45 | 0.96 | 13.55 | 1 | 10.67123 | 1.139644 |
| **2** | 12.65 | 1.14 | 8.3 | 0.7 | 7.73 | 0.85 | 11.28 | 0.87 | 12.69 | 0.94 | 13.12318 | 0.932436 |
| **3** | 11.99 | 1.05 | 7.8 | 0.95 | 8.05 | 0.65 | 12.87 | 0.98 | 13.74 | 1.02 | 14.33189 | 1.036441 |
| **4** | 13.89 | 1.33 | 8.3 | 0.51 | 9.03 | 0.91 | 10.98 | 0.89 | 12.72 | 0.83 | 12.70876 | 0.794298 |
| **5** | 13.22 | 0.97 | 8.2 | 0.97 | 6.71 | 0.45 | 14.17 | 0.84 | 14.58 | 1.13 | 13.22678 | 0.932436 |
| **6** | 12.37 | 1.14 | 9.3 | 0.7 | 7.63 | 0.96 | 13.98 | 1.04 | 13.313 | 1.35 | 13.34945 | 1.070536 |
| **7** | 13.79 | 0.86 | 8.6 | 0.57 | 8.45 | 0.48 | 12.54 | 1.23 | 14.23 | 0.97 | 12.46702 | 1.036041 |
| **8** | 13.35 | 1.031 | 9.1 | 1 | 8.16 | 0.78 | 11.87 | 0.93 | 12.06 | 0.86 | 14.36643 | 0.96897 |
| **9** | 12.81 | 1.05 | 8.9 | 1 | 7.96 | 0.94 | 13.87 | 0.96 | 13.77 | 0.89 | 14.12468 | 1.070576 |
| **10** | 13.92 | 1.35 | 7.6 | 0.81 | 8.25 | 0.56 | 12.37 | 1.12 | 12.14 | 0.86 | 13.32443 | 0.932436 |
| **Avg** | 13.245 | 1.0781 | 8.5 | 0.811 | 7.811 | 0.736 | 12.738 | 0.982 | 13.2793 | 0.985 | 13.16938 | 0.991381 |
| **Stdev** | 0.784605 | 0.168544 | 0.557773 | 0.181747688 | 0.838775 | 0.189924 | 1.074400298 | 1.13363839 | 0.810425 | 1.13363839 | 1.034652 | 1.133638 |
| **T-test** | | | 9.08E-14 | 0.000990996 | 1.19E-12 | 6.71E-05 | 0.118010201 | 0.069268689 | 0.461348 | 0.097468519 | 0.212665 | 0.104417 |

Siliques were taken from six week old plants. Mean value and standard deviation from replicates (N = 10). Level of significance was compared between the wild-type and transgenic using T-test (one-tailed, two sample with equal variance) for the statistical analysis for three replicates. Transgenics were ** significantly different from WT ($p<.01$).
